# Regulatory signatures of drought response in stress resilient *Sorghum bicolor*

**DOI:** 10.1101/2020.08.07.240580

**Authors:** Rajiv K. Parvathaneni, Indrajit Kumar, Max Braud, Philip Ozersky, Todd C. Mockler, Andrea L. Eveland

**Affiliations:** Donald Danforth Plant Science Center, St. Louis, MO 63132, USA

## Abstract

The effects of drought stress can be devastating to crop production worldwide. A grand challenge facing agriculture is development of crop varieties with improved drought resilience through breeding or biotechnology. To accelerate this, a mechanistic understanding is needed of the regulatory networks underlying drought response in crop genomes and the genetic elements that modulate them. Here, we explore the regulatory landscape of sorghum [*Sorghum bicolor* (L.) Moench] in response to controlled-environment drought stress. Sorghum is a C_4_ cereal crop with innate drought resilience. To define molecular signatures of drought response, we mapped genome-wide chromatin accessibility using an Assay for Transposase Accessible Chromatin by sequencing and analyzed parallel transcriptional profiles in drought-stressed sorghum shoot and root tissues. Drought-responsive changes in accessibility were largely in proximal promoters of differentially expressed genes and also in distal regions. Data were integrated to infer gene network connections and *cis*-regulatory modules underlying drought response and the transcription factors that control them. Inspection of pan-genomic data and phenotyping across sorghum diversity revealed variation in genomic signatures that associated with water use efficiency. Our analyses provide drought-inducible regulatory modules in sorghum that can be leveraged for fine-tuning responses to stress, mining for advantageous alleles, and translating across species.

## INTRODUCTION

Crop loss due to drought stress can be devastating. Climate shifts are predicted to become increasingly more erratic, leading to water deficits and higher temperatures that result in expansion of drought-affected arable land (Stocker et al., 2013; Varshney et al., 2018b; Gupta et al., 2020). Development of drought resilient crop varieties is necessary to help mitigate this challenge and enhance global food security (Hu and Xiong, 2014; Reynolds et al., 2016). Sorghum [*Sorghum bicolor* (L.) Moench] is the fifth most widely grown cereal crop in the world and an important staple in semiarid and arid regions of sub-Saharan Africa and south Asia (Vietmeyer et al., 1996). Because of its innate resilience to drought, heat and low nutrient inputs, sorghum presents an excellent model for studying the molecular basis for abiotic stress tolerance (Tuinstra et al., 1997; Boyles et al., 2019). Sorghum also boasts extensive phenotypic and genetic diversity (Morris et al., 2013; Lasky et al., 2015), and its adaptation to a wide range of environments provides an untapped resource for gene discovery and an ideal target for accelerated crop improvement through genomics-enabled breeding or engineering. In addition to being a staple grain crop, sorghum has emerged as an attractive system for the development of dedicated bioenergy feedstocks on marginal soils with minimal water and nutrient inputs (Mullet et al., 2014; Brenton et al., 2016). Sorghum also occupies a key evolutionary node that bridges studies in annual diploid grasses such as *Zea mays* and *Setaria*, and its close perennial and polyploid relatives, *Saccarum* and *Miscanthus*. A comprehensive understanding of the genetic factors underlying stress resilience in sorghum and the effects of genotype on phenotype, will help pinpoint targets for its improvement as well as in other cereals through comparative genomics.

Drought response in plants is a combination of cellular, morphological, and physiological components (Hu and Xiong, 2014), some of which are mirrored in other abiotic stresses that also cause dehydration, such as cold and salt stress (Nakashima et al., 2014). Plant response to dehydration at the cellular level includes production of osmoprotectants such as proline and trehalose, and stress-protecting, hydrophilic proteins like Late Embryogenesis Abundant (LEA) proteins. LEA protein synthesis promotes desiccation tolerance in seeds, but their production is also induced in vegetative tissues in response to drought (Battaglia et al., 2008; Olvera-Carrillo et al., 2011). It is suggested that regulatory modules that control *LEA* gene expression and other aspects of seed desiccation tolerance have been re-deployed to enhance drought tolerance in plants (Lamaoui et al., 2018; Pardo et al., 2020). Overexpression of *LEA* genes has been shown to confer stress tolerance in various crops (Chandra Babu et al., 2004; Duan and Cai, 2012; Amara et al., 2013), as well as through natural variation in their promoters (Xiao et al., 2007).

A major integrator of drought response in plants is the isoprenoid phytohormone abscisic acid (ABA). ABA enhances adaptation to drought and other dehydration stresses by regulating a range of physiological processes including stomatal aperture dynamics and protein storage for osmoprotection (Yoshida et al., 2015; Sah et al., 2016). ABA biosynthesis is induced under drought stress (Qin and Zeevaart, 1999; Endo et al., 2008), and its rapid local synthesis is regulated in part via hydraulic signals from the root (Christmann et al., 2007, 2013). In the leaf, ABA-induced stomatal closing allows plants to control water loss by reducing transpiration, but this also limits gas exchange for photosynthesis (Munemasa et al., 2015). Therefore, root-to-shoot coordination of ABA signaling limits above-ground biomass accumulation while enhancing water uptake from the soil. While the ABA pathway itself has been a major target of stress response mitigation, it strongly interfaces water use and photosynthetic efficiency, key traits for enhancing drought tolerance in crops (Reynolds et al., 2016; Bailey-Serres et al., 2019).

Drought tolerant genotypes tend to be associated with morphological traits like stay-green, decreased canopy leaf area, decreased stomatal density, increased leaf thickness or folding, and enhanced root system architecture; and physiological traits such as reduced transpiration rate and stomatal conductance, CO_2_ assimilation rate and canopy temperature depression (Hu and Xiong, 2014). Variation in morpho-physiological characters that contribute to abiotic stress tolerance is modulated at the molecular level by the spatiotemporal activity of transcription factors (TFs) (Joshi et al., 2016; Mittal et al., 2018). Various TF families, such as dehydration-responsive element-binding factor (DREB), basic leucine zipper (bZIP), and NAM-ATAF-CUC2 (NAC), have been implicated in regulating drought responses (Nakashima et al., 2014; Takasaki et al., 2015; Wang et al., 2019). The ABA-responsive element-binding factors (ABRE/ABFs) are bZIP TFs that drive ABA-dependent responses through positive relay of ABA perception by SNF1-related kinase 2s (SnRK2s) (Fujita et al., 2013; Yoshida et al., 2015). Numerous studies have demonstrated improved drought tolerance in crops by overexpressing or inducing stress-related TFs (Hu and Xiong, 2014; Lamaoui et al., 2018).

TFs influence target gene expression through binding of non-coding, *cis*-regulatory elements (CREs) in the genome. CREs typically occur in combinatorial arrangements in proximal promoters or distal regulators, and the coordination of chromatin accessibility and spatiotemporal expression of regulatory TFs orchestrates plant growth and development (Swift and Coruzzi, 2017; Brkljacic and Grotewold, 2017). When TF complexes bind DNA, they displace nucleosomes and the genomic region around their binding site becomes more accessible. Genome-wide maps of chromatin accessibility can therefore be used as a proxy for defining functional regions of DNA. Several methods have been used to generate accessibility maps in plant genomes including nuclease-based assays such as DNase I hypersensitivity and micrococcal nuclease (MNase) (Vera et al., 2014; Oka et al., 2017; Zhao et al., 2020), and an Assay for Transposase Accessible Chromatin (ATAC), which uses a Tn5 transposase that inserts into accessible regions of DNA (Buenrostro et al., 2015). Studies based on these methods have revealed dynamic shifts in chromatin accessibility during development or in response to stimuli (Sullivan et al., 2014; Reynoso et al., 2019), as well as conserved and unique regulatory signatures across species (Maher et al., 2018; Lu et al., 2019; Han et al., 2020). Accessible chromatin has been associated with heritable variation (Rodgers-Melnick et al., 2016; Parvathaneni et al., 2020) and can guide construction of gene regulatory networks through inferences of TF-CRE interactions (Sullivan et al., 2014).

A knowledgebase of the genes and regulatory sequences that underlie stress responses in plants will enable predictions on gene regulation that can help improve crops. In this study, we explore the drought-responsive genome in sorghum and define gene regulatory modules that potentially underlie its resilience. Using RNA-seq-based transcriptome analyses coupled with ATAC-seq from the same sample preps, we annotated drought-related signatures in developing leaf and crown root tissues in response to a controlled-environment drought stress. Using a co-expression gene regulatory framework across tissues, we inferred core TF-DNA interactions and combinatorial regulation of downstream target genes in response to stress. Leveraging a pan-genomic resource and high-throughput phenotyping across sorghum diversity, we identified variation in core drought-inducible signatures that associated with water use efficiency (WUE). Our results provide a resource for elucidating molecular responses to drought in sorghum and a toolkit of drought-inducible regulatory sequences for engineering improved crops.

## RESULTS

### Genome-wide dynamics of chromatin accessibility in response to drought

To define genetic elements that underlie drought response in sorghum, including those implicated in drought resilience, we profiled the gene regulatory space in shoot and root tissues of young sorghum plants subjected to a controlled drought treatment, and well-watered (WW) controls. Sorghum plants of the reference genotype BTx623 were grown in a controlled-environment chamber and watered to 100 percent Soil Moisture Capacity (SMC) until 17 Days After Sowing (DAS). At this time, plants in the water stressed (WS) group were gradually brought down to 25 percent SMC by 21 DAS, replacing water lost by transpiration, and held for ~48 hours before destructive sampling (Figure 1A). At 23 DAS, plants in both WW and WS groups were sampled as follows: the emerging leaf was dissected out and sectioned across a developmental gradient (Li et al., 2010), the inner developing leaves within the emerging leaf whorl were sampled, and the crown roots sampled after root washing (Figure 1B). All samples were collected for transcriptome profiling using RNA-seq.

**Figure 1.**
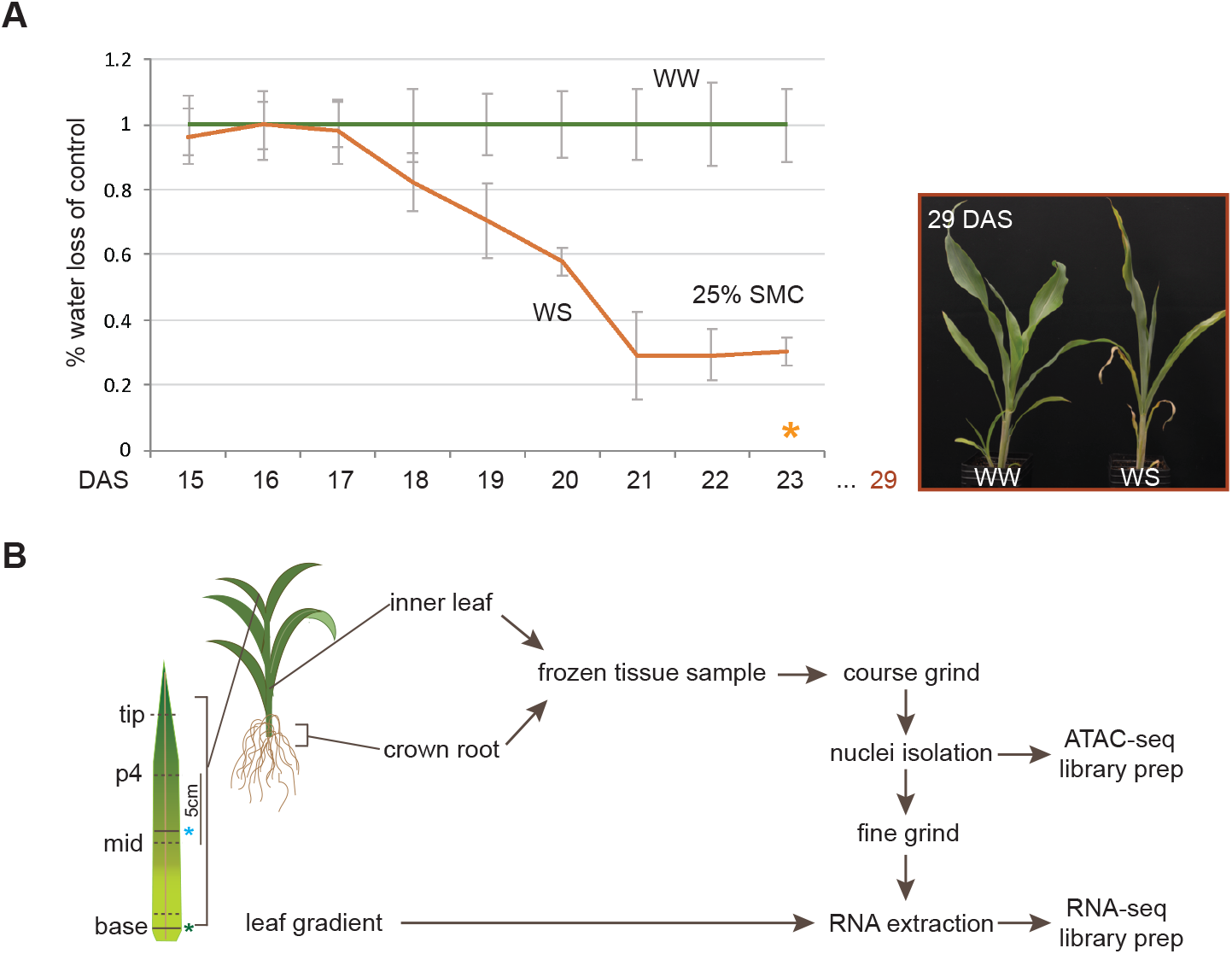
Experimental design for annotating regulatory components of drought response in sorghum seedlings. **A)** Controlled watering strategy imposed on sorghum seedlings from well-watered (WW) and water stressed (WS) groups in a controlled-environment chamber. Plants were sampled at 23 DAS (48 hrs after hitting 25 SMC; orange star). Image of WW and WS plants taken at 29 DAS shows post drought treatment effects on plant morphology. **B)** Sampling strategy for collecting RNA-seq and ATAC-seq data across tissues from the same plant. Four 1 cm sections were sampled along the gradient of the emerging leaf. Green star = ligule; blue star = point at which leaf emerges from the whorl; p4 = 4 cm above the point from which leaf emerges; mid - 1 cm below point at which leaf emerges.

For the inner developing leaf and crown root samples, we also profiled chromatin accessibility using ATAC-seq from the same exact sample used for RNA-seq. Each biological replicate represented a single plant, and we analyzed three biological replicates in each group. Each individual leaf or root sample was lightly ground in liquid nitrogen and then divided in half. Nuclei were isolated from one half and used immediately for ATAC-seq library preparation (Supplemental Figure 1). The other half was ground further for subsequent RNA extraction and RNA-seq library construction. ATAC-seq libraries were sequenced and mapped to the sorghum reference genome v3 (phytozome.jgi.doe.gov; Supplemental Table 1). Overall, we achieved higher coverage for the leaf datasets. Visual scans of the ATAC-seq data on a JBrowse viewer and correlation analysis revealed high concordance across biological replicates genome-wide (Figure 2A; Supplemental Figure 1). Regions of significant read pile-ups were called as ‘peaks’ using the algorithm HOMER (Heinz et al., 2010), and serve as proxies for chromatin accessible regions (Supplemental Data Set 1). Most of the accessible genome was shared across tissues and treatment groups (Figure 2B).

**Figure 2.**
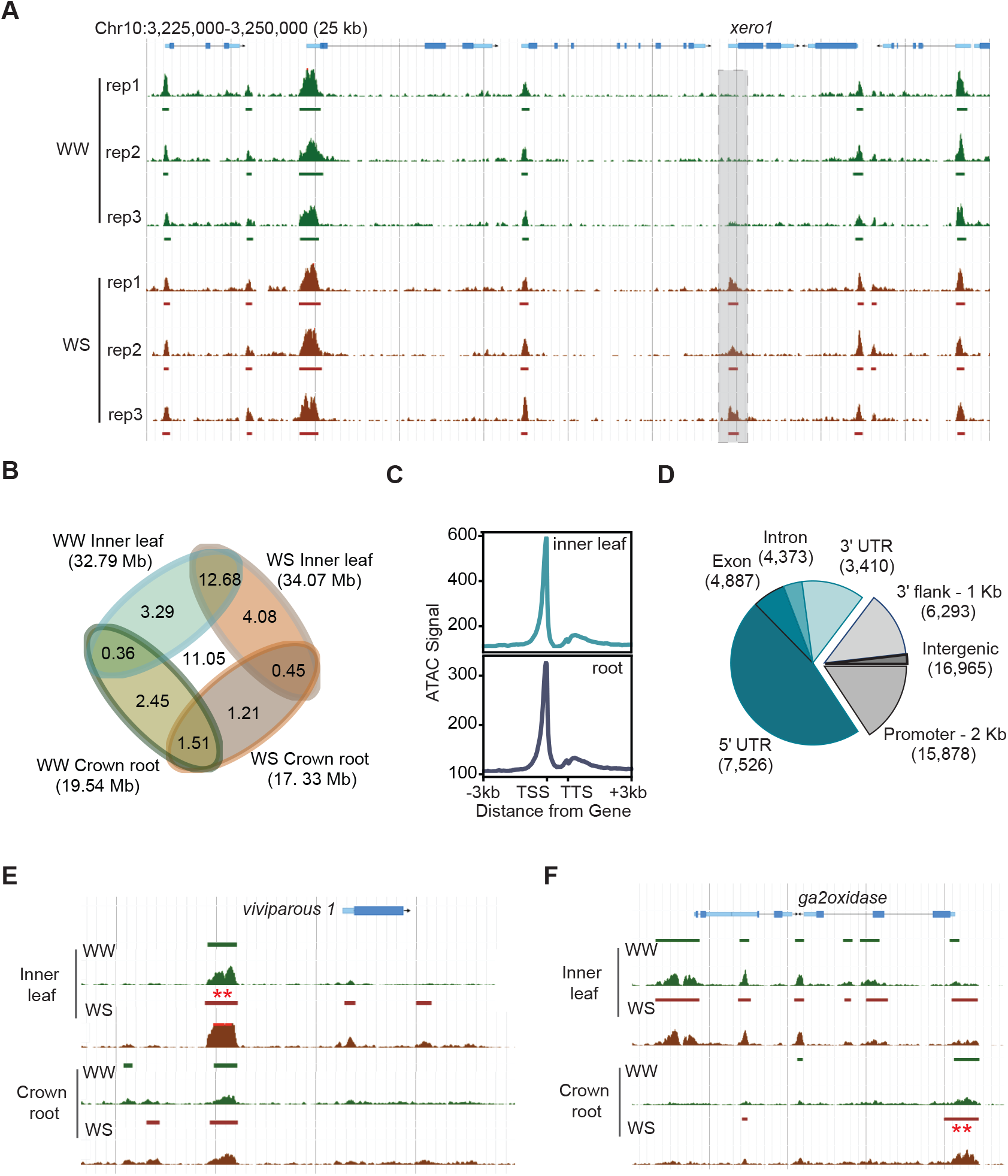
Genome-wide signatures of chromatin accessibility in sorghum shoot and root tissue in response to drought. **A)** Profiles of chromatin accessibility by ATAC-seq in inner developing leaf tissue are shown across biological replicates and in control (WW) and drought stress (WS) conditions for a genomic region in the sorghum genome. The *xerico* (*xero*) *1* gene is highlighted with a drought-specific signature in its promoter. **B)** Overlap of genome accessible space across tissues and treatments. ~83% of the accessible genome was shared between drought and control treatments (87% in leaf and 78% in root) and ~78% was shared between tissues (75% in control and 81% in drought). **C)** Read coverage of the ATAC-seq across the consensus gene model in inner leaf and root tissue. A window of 2kb upstream and downstream of the consensus gene model was used. **D)** Distribution of the control inner leaf and root ATAC peaks across the genomic features. **E)** A leaf-specific DAR is shown 3 kb upstream of the *vp1* ortholog. **F)** Root- and leaf-specific chromatin accessibility shown around a gene encoding *GA-2 oxidase*.

We defined a set of ‘unique peak’ regions of accessibility across both tissues and treatments that showed consistency across biological replicates (see Methods; Supplemental Table 2, Supplemental Data Set 2). Coverage of ATAC-seq reads in relation to protein-coding gene models showed strong enrichment of signal in the proximal promoter regions, and to a lesser extent at the 3’ ends (Figure 2C). Distribution of unique peaks across genic and intergenic features showed that indeed, the majority of these accessible regions were located within the proximal promoters and 5’ UTRs of genes (Figure 2D; Supplemental Table 3). In addition, a substantial portion (29%) of accessible peaks were categorized as intergenic (excluding space within 2 kb up- and 1 kb down-stream of annotated gene models).

To define regions of the genome that showed differential accessibility in response to drought, we adapted a pipeline from (Sullivan et al., 2019). High-confidence ATAC peaks from both treatment groups were merged into a set of ‘union peaks’, and this was done separately for leaf and root (see Methods). We used a statistical test based on differences in numbers of ATAC transposition sites for each union peak between WW and WS plants to determine Differentially Accessible Regions (DARs). Across the sorghum genome, we identified 6,496 and 2,275 drought-responsive DARs (median size ~443 bp) in developing inner leaf and crown root, respectively, and identified the closest gene model(s) within 10 kb to each DAR (Supplemental Data Set 3; Supplemental Table 3). This allowed us to link intergenic DARs to proximal genes and to investigate gene-associated DARs across tissues. For example, a DAR positioned 3 kb upstream of the sorghum ortholog of maize *viviparous 1* (*vp1*; Sobic.009G221400) showed increased accessibility under drought in the leaf (Figure 3E). A DAR in the promoter region of a gene encoding a *GA-2 oxidase* (Sobic.004G222500) showed a root-specific drought response and in the gene itself there is a root-specific DAR. The gene just proximal to it (Sobic.004G222400) showed leaf-specific accessibility (Figure 3F).

**Figure 3.**
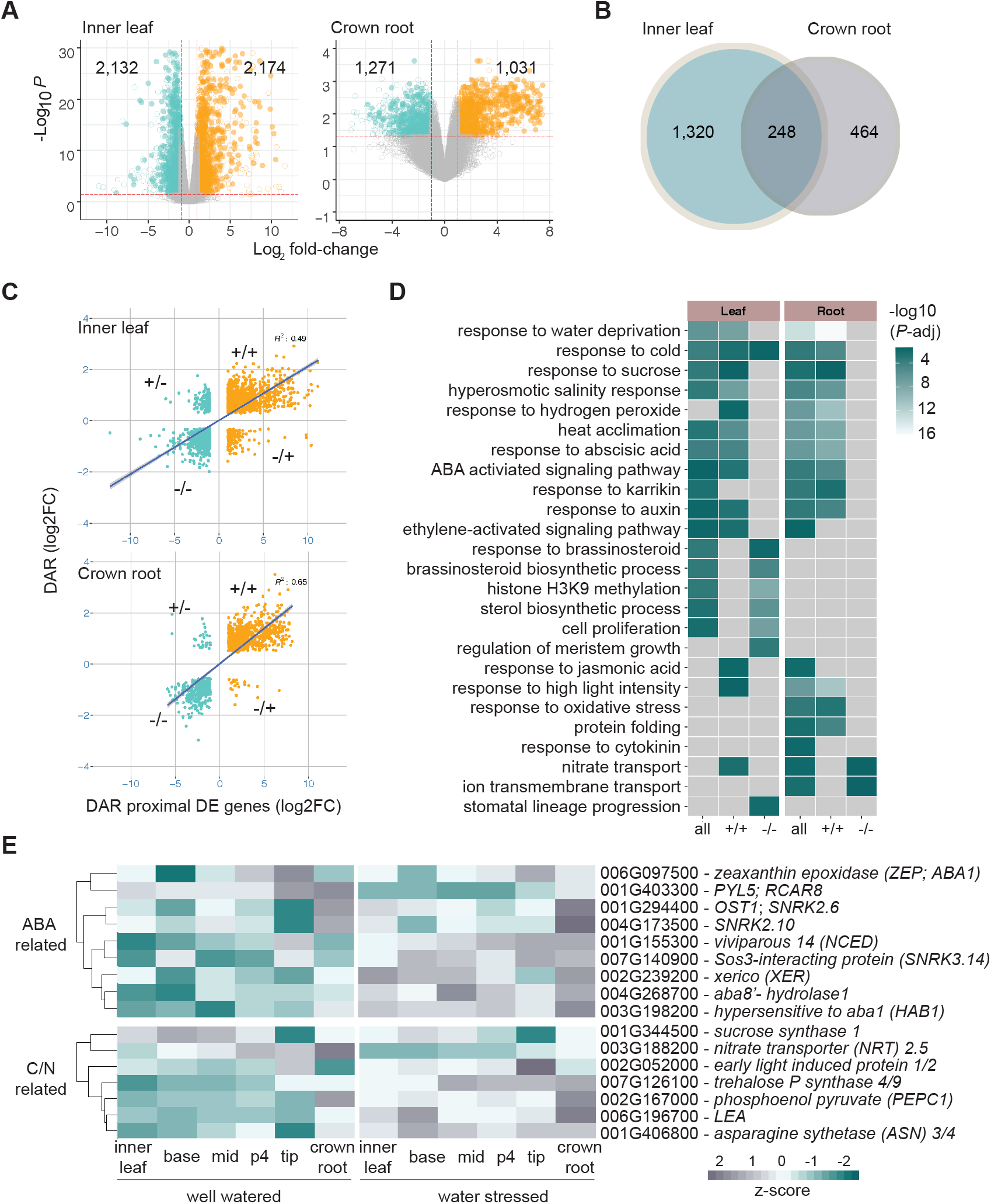
Drought-responsive transcriptional changes are associated with differential chromatin accessibility. **A)** Volcano plots show the distribution of DE genes in the developing leaf and crown root tissues in response to drought stress (orange circles = up-regulated genes; light blue circles = down-regulated genes; filled circles represent DAR-proximal DE genes). **B)** Differentially regulated genes that are common and distinct between the inner leaf and crown root in response to drought. **C)** Scatterplots show strong correlations between direction of DAR accessibility (more accessible = +; less accessible = -) and changes in expression of corresponding gene (increased expression = +; decreased expression = -). **D)** Functional classes of genes based on GO enrichment analyses for subsets of differentially regulated genes. **E)** Expression profiles across a developing leaf gradient and in crown roots for a subset of drought-responsive, DAR-proximal genes that are associated with physiological responses to drought.

### Drought-induced gene regulation links hormone, stress, and physiology pathways in sorghum

We also profiled transcriptional changes in response to drought using RNA-seq in the exact same samples processed for ATAC-seq. In the inner leaf, 4,307 genes were differentially expressed (DE) in response to drought (FDR ≤ 0.05; ≥ 2-fold change), and 2,303 in crown roots (Figure 3A; Supplemental Figure 2; Supplemental Data Set 4). Overall, gene expression differences in response to drought were larger in the developing leaf compared to the crown root (Figure 3A). In addition, genes that were up-regulated in the leaf tended to show larger differences in expression, which coincided with enhanced chromatin accessibility at their promoters, and this was not observed for down-regulated genes (Supplemental Figure 3). Among the DE genes, 1,568 (36%) in inner leaf and 712 (31%) in crown root were associated with DARs, and 248 of these were shared between both tissues (Figure 3B; Supplemental Data Set 3). These genes are hereinafter referred to as differentially regulated. We classified differentially regulated genes as having correlated or anti-correlated expression values relative to the direction of accessibility change in the associated DAR. For example, is increased (+) or decreased (−) gene expression associated with more (+) or less (−) DAR accessibility in response to drought? We observed a strong positive correlation among gene expression and accessibility, with a higher number of genes positively correlated with DARs than anti-correlated (Figure 3C). This potentially suggests more prominent examples of TF activator activity compared to repressor activity.

Gene Ontologies (GO) were used to test for overrepresentation of functional classes among differentially regulated genes. In both developing leaf and crown root, there was significant enrichment of genes associated with various, but related, stress responses (e.g., water deprivation, cold, salinity), sugar metabolism, and several hormone response pathways, including ABA (Figure 3D; Supplemental Data Set 5). Certain functional categories showed tissue-specific enrichment and there were also differences among genes showing up- and down-regulation in leaf and root tissues (Figure 3D). For example, DE genes specifically up-regulated in the crown root were enriched for functional categories related to oxidative stress and protein folding, while down-regulated genes in the developing leaf were enriched for functional categories associated with developmental progression; e.g., ‘regulation of meristem growth’ (GO:0010075) and ‘stomatal lineage progression’ (GO:0010440). The latter being consistent with repression of developmental processes in response to stress, and perhaps reduction of stomatal patterning in early leaf development. Most genes related to hormone synthesis and signaling were up-regulated and associated with increased chromatin accessibility (+/+) in leaf and root, however biosynthesis and response of brassinosteroids (BRs) showed dampened regulation (−/−) in the leaf. Response to cytokinin was enriched only among differentially regulated genes in the root. Interestingly, ‘nitrate transport’ (GO:0010167) was overrepresented among genes that were up-regulated in leaf, but down-regulated in root (Figure 3D).

In addition to transcriptional changes in the developing inner leaf, we also profiled gene expression along the developmental gradient of the emerging leaf in response to drought. RNA-seq was performed on four distinct sections from base to tip, which capture distinct zones of developmental and metabolic transitions in grass leaves (Figure 1A; Supplemental Data Set 6). The base leaf section, closest in progression to the developing inner leaf, is where developmental decisions are still occurring, while C_4_ photosynthesis and associated metabolism progresses towards the leaf tip (Li et al., 2010). As was shown in maize, *Setaria*, and rice, suites of TFs were differentially regulated along the developmental timeline of the leaf (Wang et al., 2014) as well as hormone responses (Supplemental Data Set 6). We also observed dynamic expression profiles in response to drought for many differentially regulated genes, including those with known functions in ABA synthesis and signaling as well as related to carbon (C)-nitrogen (N) metabolism, allocation and use and osmoprotection (Figure 3E).

### Drought-responsive DARs are enriched for binding sites of stress-associated TFs

To infer putative TF binding sites within DARs proximal to drought-responsive genes, we tested for enrichment of CREs. DARs were grouped based on their response to drought (more (+) or less (−) accessible), and the correlation of this response with that of the proximal genes (expression increase (+) or decrease (−)). An enrichment test for *de novo* motifs was performed on these subsets of DARs and Position Weight Matrices (PWMs) for enriched sequences were compared to known PWMs in the JASPAR plant database. Best matches to experimentally validated PWMs from plants were reported (Supplemental Data Set 7), which include binding sites for several TF families previously implicated in abiotic stress response; e.g., ABRE, DREB, and NAC.

We compared *de novo* motifs that were discovered between leaf and root, and among up- and down-regulated gene sets using STAMP (Mahony and Benos, 2007). This revealed tissue-specific preferences for certain motifs as well as motifs that were preferentially found in DARs associated with drought induced or repressed genes (Figure 4A; Supplemental Data Set 7). Among elements enriched in (+/+) DARs in both tissues were PWMs related to the bZIP HY5/AREB1 and the AP2/ERF CYTOKININ RESPONSE FACTOR 4 (CRF4) TF motifs (Figure 4A, Supplemental Data Set 7). A putative BRAZZINOLE-RESISTANT 1 (BZR1) binding site was also enriched in (+/+) DARs, even though (−/−) DE genes were enriched for BR-related functional categories (Figure 3D). ABRE-like sequences (GMCACGY; E-value = 2.3e-41) were among the most enriched in (+/+) DARs in both inner leaf and crown root (Figure 4A). Expression of ABA-induced genes is typically driven by the presence of multiple ABA-responsive elements (ABRE) motifs or a combination of ABRE with other elements in the promoter (Yoshida et al., 2010; Shen et al., 1996; Hobo et al., 1999). A well-characterized aspect of this regulation in promoting desiccation tolerance is coordinated regulation of ABA responsive genes by the bZIP TF ABI5 (ABRE) and B3 TF ABI3 (orthologous to maize *vp1*). The (−/−) DARs showed enrichment for bHLH binding sites, consistent with roles for these TFs in regulation of cell division and identity, for example, in stomatal patterning. Certain motifs showed tissue specificity such as those related to ERF4- and ABI3-related PWMs, which were largely enriched in inner leaf-specific DARs (Figure 4A).

**Figure 4.**
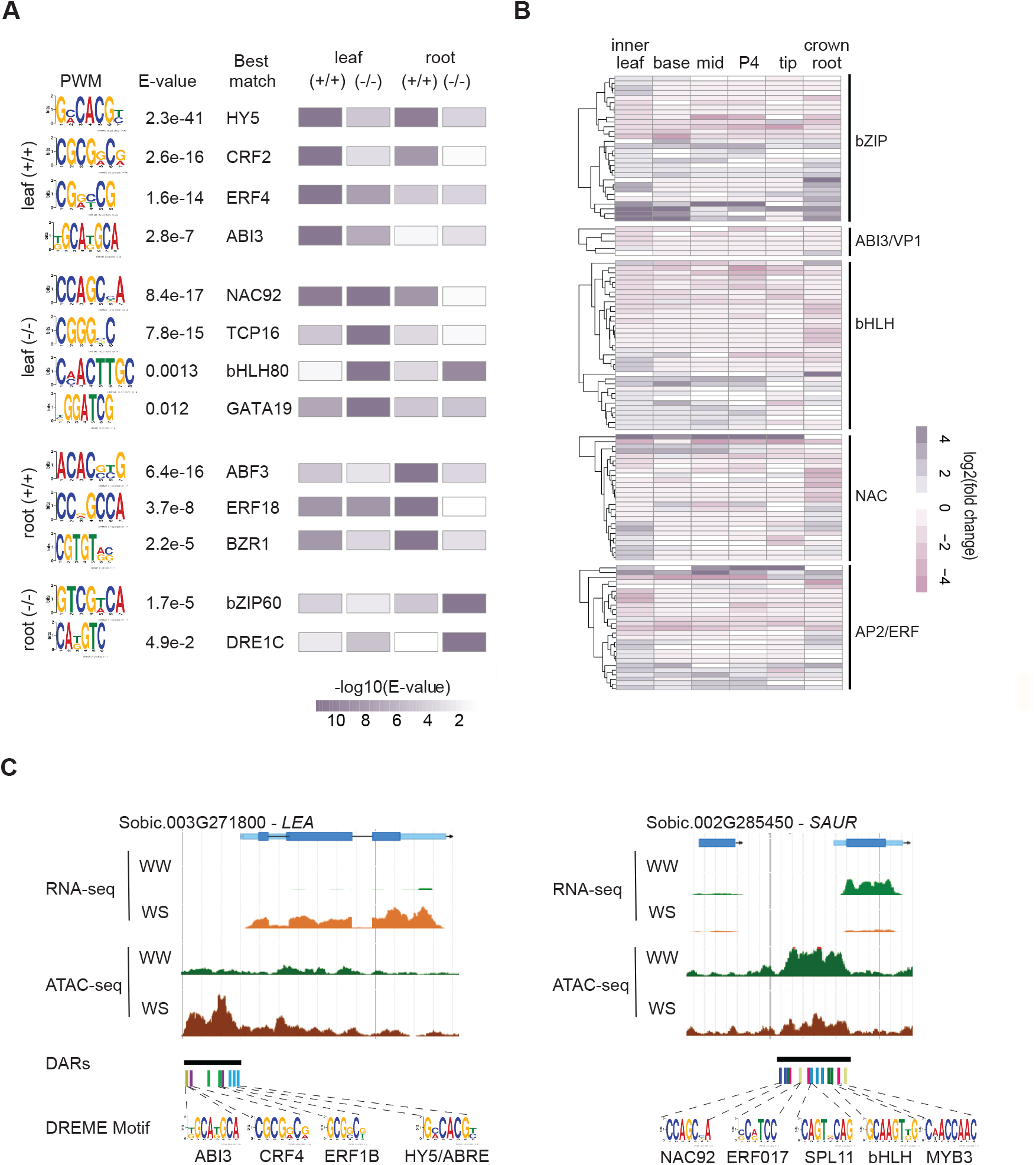
Mapping CREs underlying drought-responsive DARs. **A)** *De novo* motifs enriched in DARs that correlate with expression of proximal genes were compared based on their relative enrichment in each category (+/+ or −/−) or between leaf and root tissues. PWMs were compared with those in JASPAR Plant DB to identify top matches. The enrichment scores for each PWM were compared pairwise to motifs found in all other categories/tissues to determine a similarity value (E-value). **B)** Expression differences of TFs associated with certain enriched motifs were plotted based on their log fold change differences between WW and WS across the inner leaf, developing leaf gradient, and crown root samples. **C)** Jbrowse views showing DARs in the promoters of up-regulated *LEA* and down-regulated *SAUR* genes and presence of enriched *de novo* motifs. Presence of multiple motifs suggest combinatorial regulation by multiple TFs.

Differential accessibility and expression of target genes are expected downstream of drought-responsive transcriptional activity. We surveyed DE genes for TFs associated with the DAR-enriched PWMs. We annotated 174 DE TFs within these classes in inner leaf and/or crown root (Supplemental Data Set 8), including bZIP (30), B3 (6), bHLH (35), ERF/DREB (27), and NAC (26) TFs. These showed dynamic expression patterns across leaf and root tissues and in response to drought (Figure 4B, Supplemental Data Set 8). Consistent with bZIP PWMs being enriched in (+/+) DARs, the majority of DE bZIP TFs (70%) were up-regulated in leaf and root tissues in response to drought stress. In the case of bHLH (enriched in (−/−) DARS), DE TFs were approximately half up- or down-regulated (Supplemental Data Set 8).

Since TFs act in a combinatorial manner to modulate the stress response, we can leverage their shared expression profiles and CREs that co-occur within DARs to infer TF-DNA interactions. We used the enriched set of PWMs to scan DARs associated with DE genes to define combinations of CREs that may provide insights into the TFs that modulate them in response to drought. For example, a gene encoding a LEA protein (Sobic.003G271800) was up-regulated in response to drought and its accessible promoter harbored several CREs including CRF4 and HY5/ABRE that were enriched in leaf (+/+) DARs (Figure 4C). Alternatively, a gene encoding a small auxin up-regulated RNA (SAUR) was down-regulated in response to drought and its promoter included several CREs found associated with (−/−) DARs, such as bHLH (Figure 4C).

### Gene network analysis resolves drought-associated regulatory modules in sorghum

We expect that genes with highly similar expression profiles across tissues and treatments could potentially work together in a given pathway, and/or be regulated by a common set of TFs. To help resolve TF-DNA interactions in response to drought, we used a weighted gene co-expression network analysis (WGCNA) coupled with a random forest classifier to construct a gene regulatory network based on the collective RNA-seq data across WW and WS samples. We used the WGCNA algorithm (Langfelder and Horvath, 2008) to generate a co-expression network that included 22,847 nodes (genes) in 23 distinct co-expression modules (Supplemental Figure 4). Module eigengenes (ME; the expression pattern that best fits an individual module across all samples) were evaluated for their significant associations with specific tissue types and the drought treatment (Figure 5A). Certain MEs showed tissue-specific expression patterns; e.g., MElightcyan showed a strong association with specificity to the inner developing leaf and MEroyalblue with crown roots (Figure 5A; Supplemental Figure 5).

**Figure 5.**
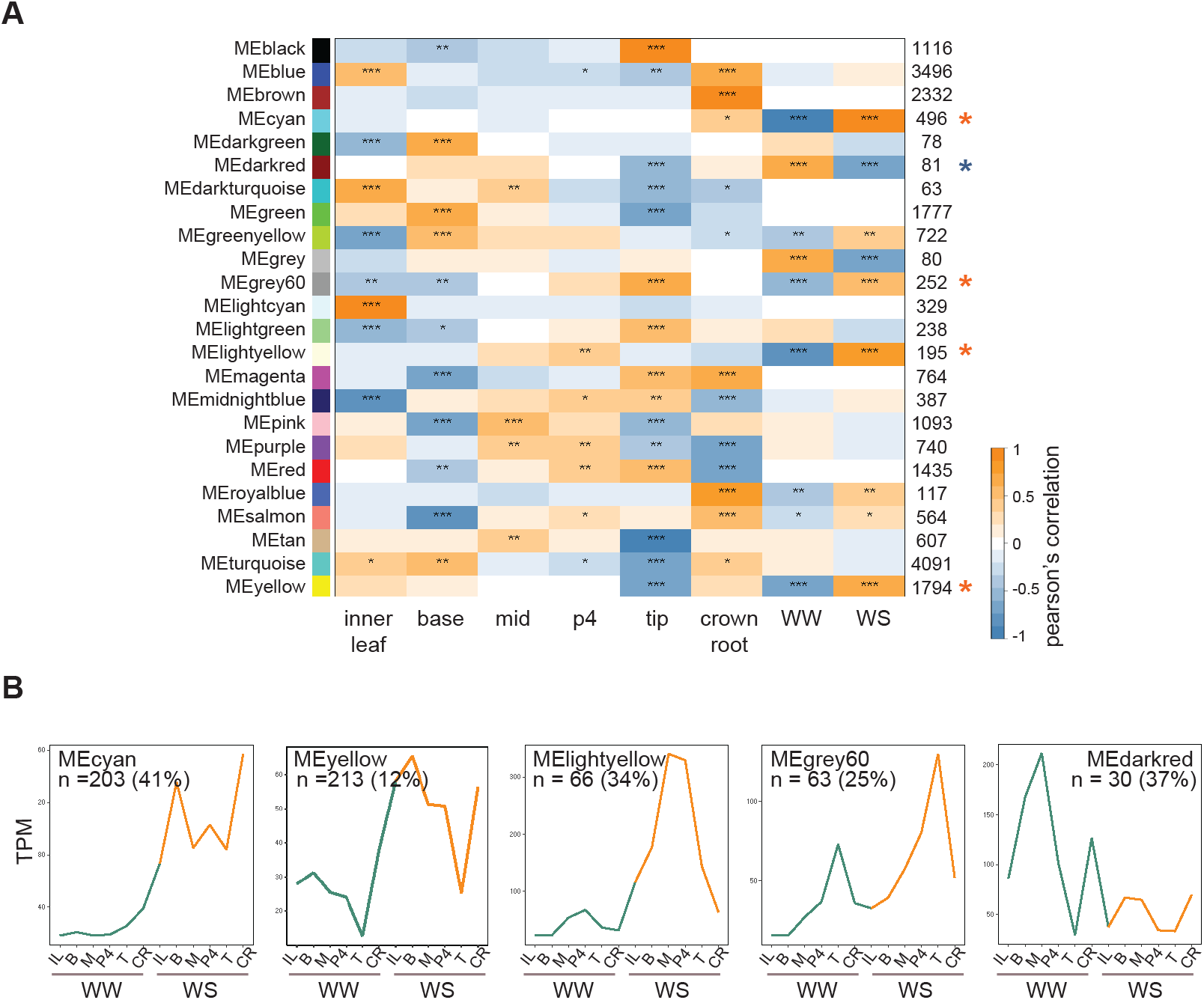
Gene network analysis identifies drought-associated co-expression modules. **A)** The heatmap represents the WGCNA eigengene module associations with specific tissues and/or drought treatment in sorghum. Network modules are represented and named with different colors based on the WGCNA default modules annotation. The number of genes co-expressed in each module are indicated to the right. Student asymptotic *p*-values for the eigengene module association are indicated: ***, *p*-value < 0.001; **, *p*-value < 0.01; *, *p*-value < 0.05. **B)** Trajectories of module eigengenes for five drought-associated modules across the WW and WS leaf and crown root samples. *n* = number of differentially regulated genes (DE and associated with a proximal DAR) in each module.

Several modules showed strong associations with the drought response. Among these, MEcyan, MElightyellow, MEyellow and MEgrey60, showed positive correlations with drought response (R^2^ > 0.50) and MEdarkred showed a strong negative correlation (R^2^ = − 0.67; Figure 5A). The ME profiles among positively regulated modules (MEcyan, MEyellow and MElightyellow) were closely related but showed distinct features (Figure 5B; Supplemental Figure 5). A relatively high proportion of genes within cyan and lightyellow modules, 41 and 34 percent, respectively, were differentially regulated in response to drought in either inner leaf or crown root. Among genes within these modules we observed overrepresentation of many functional categories related to various types of dehydration stress, ABA signaling, and osmotic adjustment (Supplemental Data Set 9).

We also integrated the co-expression network with information derived from regulatory interactions among TFs and their putative targets using the GENIE3 algorithm (Huynh-Thu et al., 2010). We annotated 1,277 TFs in the network based on PlantTFDB (Jin et al., 2017) and used their network trajectories to identify connections with potential target genes. There were 179 TFs among the five drought-associated modules and we focused on TF-target interactions within these modules. This enabled us to identify suites of genes that were co-expressed in response to drought, and to predict potential TF-DNA interactions based on motif scans of DARs within promoters of co-expressed genes. To do this, we performed motif enrichment analyses within DARs associated with DE genes in the five drought modules, and then scanned their differentially accessible promoters for enriched motifs to determine co-occurrences of CREs across these drought-associated regulatory regions (Supplemental Data Set 7). This provided strength for predictions on TF-DNA interactions based on the GRN.

Of the TFs associated with the five drought modules, 84 were DE in response to drought and 37 were predicted to be network hubs across these modules. Four of these TFs were identified as putative AREB/ABFs based on homology. While the regulatory networks associated with action of ABA-induced ABF TFs have been well-studied in Arabidopsis (Song et al., 2016), relatively little is known about their intricacies in grasses. We identified four enriched motifs from the DREME analysis that contained the “ACGT” core ABF binding motif (ABRE) sequences, which were further used for presence/absence screening in DARs of putative target genes. For example, we focused on two ABFs that showed strong connectivity in the network, and whose targets were highly enriched in drought-related GO terms (Supplemental Data Set 10); Sobic.010G081800 (SbAREB1/ABF2) and Sobic.007G155900 (SbABF3). We filtered their predicted direct targets by whether they were also DE, DAR proximal and contained enriched ABF motifs (Supplemental Data Set 11). This resulted in 51 SbABF2 and 38 SbABF3 targets, of which 23 were shared. Predicted ABF targets include several known players in drought response such as PP2C, ABI-binding, SNRK3.14, SNF1, heat shock proteins, and ABI1-like (Figure 6). All but three of the combined ABF targets were up-regulated in response to drought.

**Figure 6.**
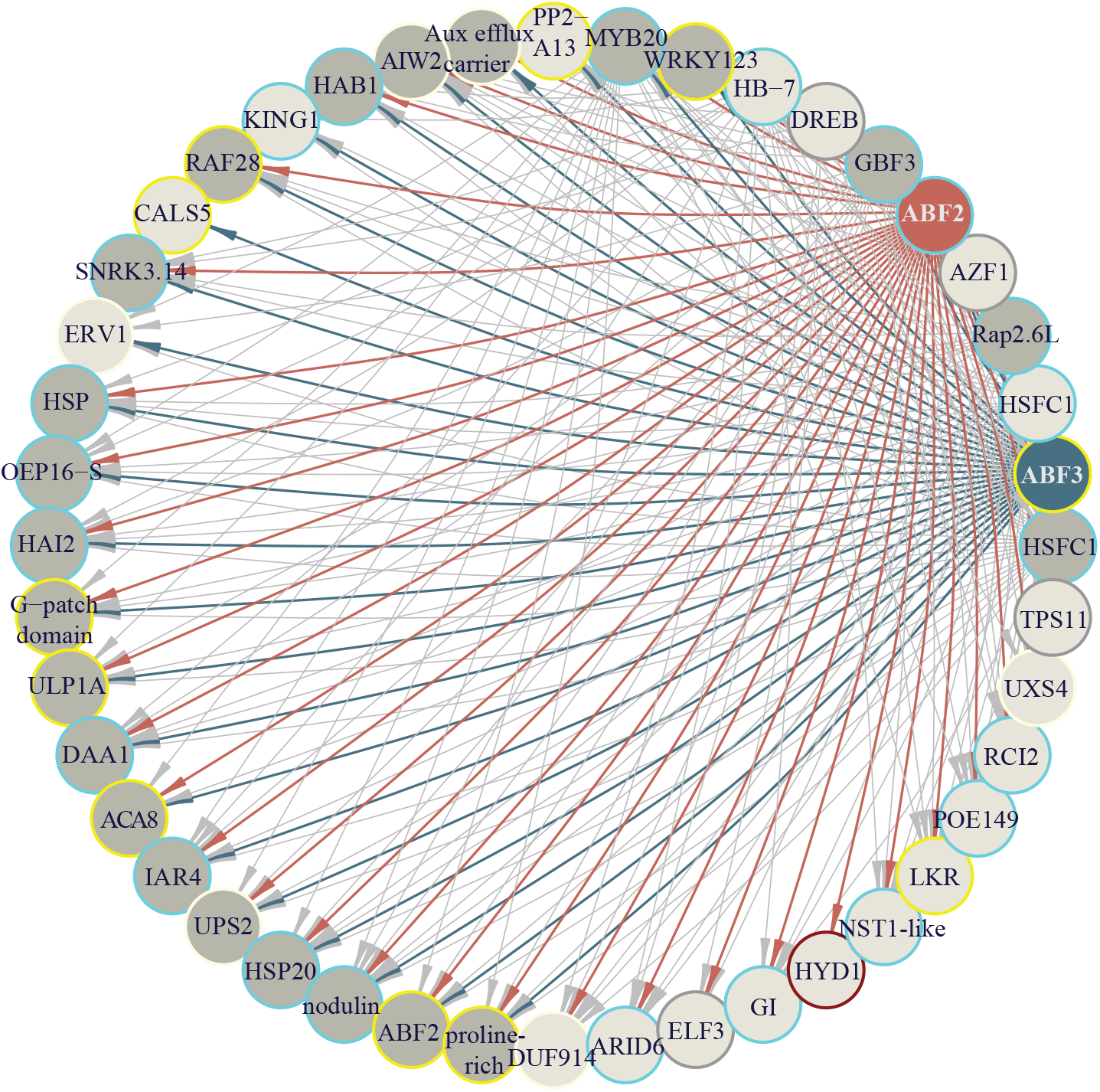
Sub-network of two hub ABF TFs reveals overlapping and unique target genes. Sorghum orthologs of AREB1/ABF2 (red) and ABF3 (blue) were identified as hub TFs in the cyan and yellow modules, respectively. Based on GRN predictions, direct target genes of these ABFs within the five drought-associated modules were classified as high confidence if they had a DAR within their upstream promoter (2 kb) and curated ABF binding motifs within those DARs. Nodes (genes) are represented as circles with edges connecting to their predicted targets. Targets common between ABF2 and 3 are colored dark grey and unique targets are light grey. The outline color of each node indicates the module identity from the WGCNA. Additional TF-target gene connections aside from ABFs shown are based on GRN predictions only.

### Sequence variation in the drought-responsive genome associates with WUE

Our analyses provide a catalog of drought-induced regulatory features, including hub TFs, their target genes and combinatorial binding sites. We expect that sequence variation in these features across sorghum diversity contributes to regulatory function in response to drought, and can be leveraged to identify beneficial alleles for breeding enhanced stress resilience. To survey sequence diversity associated with candidate genes and regulatory elements underlying drought response, we leveraged a sorghum pan-genome resource (https://terraref.org), which catalogs tens of millions of sequence variants including single nucleotide polymorphisms (SNPs), indels and presence/absence variation (PAV) based on resequencing data from 363 diverse lines. We also phenotyped 380 lines from the Bioenergy Association Panel (BAP; (Brenton et al., 2016) in a controlled-environment experiment using a LemnaTec scanalyzer to image 2,280 plants in well-watered (WW; 100% SMC) and controlled drought (WS; 30% SMC) conditions over a two week time course (Supplemental Figure 6). Water use efficiency (WUE) for each line was calculated daily based on a model that incorporates pixel area and water used (Methods; Supplemental Data Set 12) (Fahlgren et al., 2015). Of the 380 phenotyped lines, 322 were among the resequenced lines in the pan-genome resource. These lines were binned into eight distinct subpopulations (subpops) based on a population structure analysis. We observed subpop-associated trends in the median WUE across the drought response period (15-21 days after planting); e.g., subpop 2 showed overall higher WUE compared to the others, and subpop 3 showed the lowest WUE (Figure 7A; Supplemental Figure 7).

**Figure 7.**
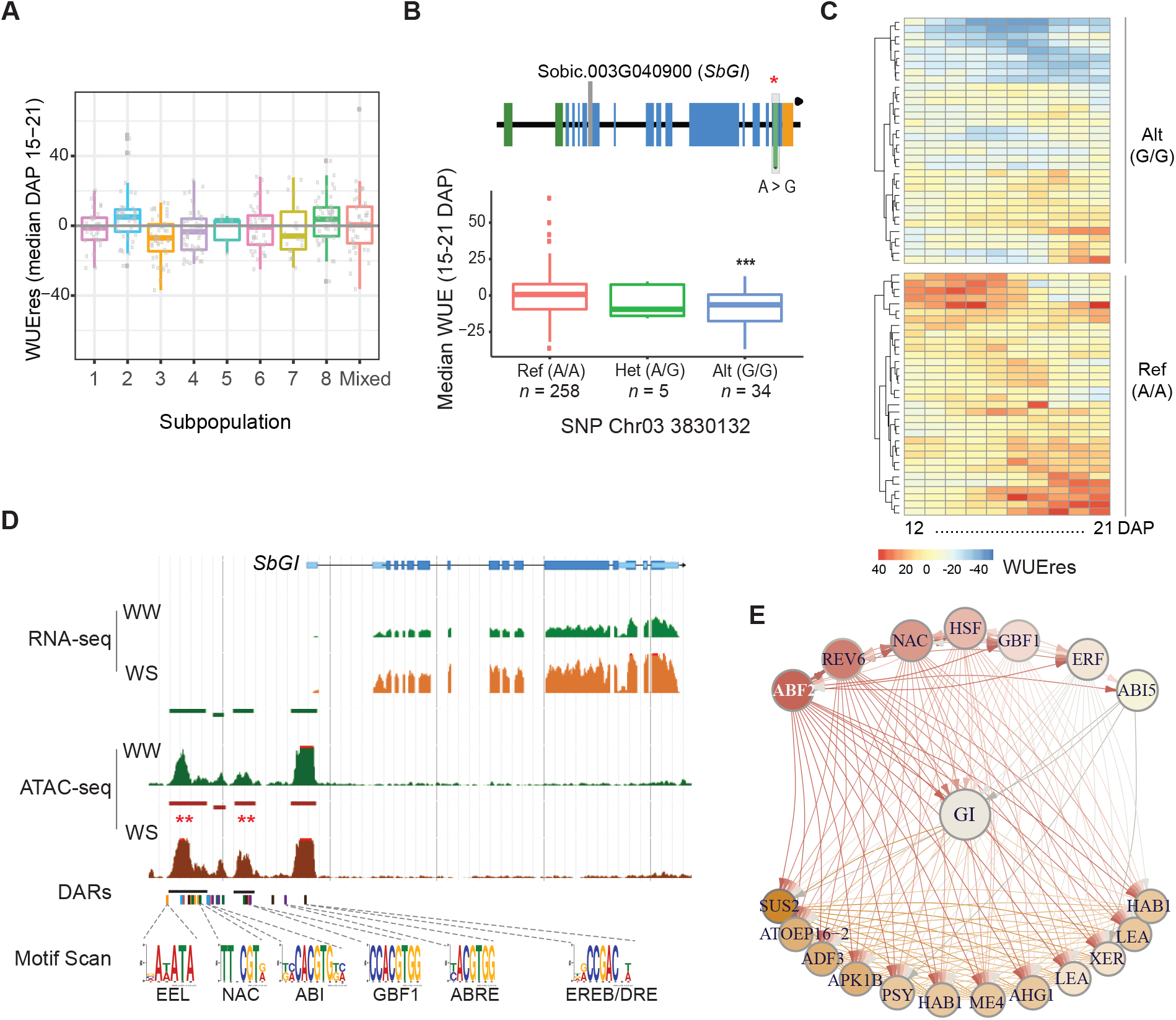
Pan-genomic analysis of drought-responsive loci reveals novel variation associated with WUE. **A)** Box plots show variation in WUE by subpopulation in response to drought stress. Analysis is based on median WUE across 7 days (15-21 days after planting) of the stress response. **B)** The *SbGI* locus with variation based on pan-genome analysis (above) shows a SNP (red star/grey box) with significant association with decreased WUE (below). The box plots show variation in WUE for lines with the Ref allele and with the Alt allele. **C)** Heatmaps show dynamic WUE residuals across 10 days of phenotyping in response to drought stress for 34 sorghum lines with the Alt *SbGI* allele compared to a random subset of 34 lines with the Ref allele. **D)** Genome browser view of the *SbGI* locus in developing leaf tissue shows increased expression in response to drought stress (RNA-seq) and differential chromatin accessibility (DARs, red **). Several motifs associated with predicted upstream TFs based on GENIE3 analysis are indicated in the promoter region. **E)** A SbGI sub-network with direct up-stream regulators as predicted by GENIE3 and a subset of nearest neighbors identified by WGCNA that were differentially regulated in response to drought. Aside from ABF2, predicted upstream regulators include: REV6 (Sobic.010G223700), NAC (Sobic.008G021800), HSF (Sobic.004G101400), GBF1 (Sobic.009G237600), ERF (Sobic.002G212000), ABI5 (Sobic.003G332200). The full list of nodes in this SbGI sub-network is provided in Supplemental Data Set 14.

To determine whether variation in core drought network components was associated with WUE, we looked at the differentially regulated targets of the ABF TFs (Figure 6; Supplemental Data Set 11) and the PAV and SNPs within them. Six out of the 70 unique ABF gene targets showed extensive PAV across subpops (i.e., absent in at least 40% of genotypes; Supplemental Figure 8A; Supplemental Data Set 13). Interestingly, three of these were not associated with any functional annotation (Sobic.006G007300, Sobic.006G007650 and Sobic.001G342800). Another encoding a Ubiquitin-like-specific protease 1a (ULP1a; Sobic.001G028900) showed PAV in most subpops (except subpop 1) and was largely absent (~80%) in subpops 3 and 6, which tended to have lower WUE (Supplemental Figure 8B). In Arabidopsis, ULP1a acts in deSUMOlyation of target proteins in the BR pathway in response to stress (Srivastava et al., 2020). Modification of proteins by a small ubiquitin-related modifier (SUMO) is a mechanism by which plants can rapidly integrate growth with response to environmental cues, for example in response to drought (Benlloch and Lois, 2018) or salt (Srivastava et al., 2020) stresses. This gene was also highly polymorphic among lines as well.

Several ABF target genes were polymorphic among lines with SNPs of strong predicted effect size, many of which tended to be subpop-specific or subpop-dominant (Supplemental Figure 9). To test whether 34 moderate-to-high effect size SNPs that were strongly associated with subpops 2 or 3 (higher and lower WUE, respectively) showed significant associations with plant WUE, we used two different statistical approaches, the Tukey HSD multiple pairwise comparison test and a Student’s t-test on 100,000 random samples of the WUE values. Based on at least one of these statistical tests, we identified four SNPs in differentially regulated ABF target genes that were significantly associated with WUE, and polymorphisms largely exclusive to subpop 3. Among these, significant associations were found within a LRR and NB-ARC domain containing gene (Sobic.002G104400), a gene encoding a mitochondrial inner membrane translocase (Sobic.003G144500), and a Ser/Thr protein kinase (HT1; Sobic.007G078300) (Supplemental Figure 9; Supplemental Data Set 13).

Variation in a SNP of predicted moderate effect size (A>G; p.Asn1095Asp) in the last exon of the sorghum *GIGANTEA* ortholog (*SbGI;* Sobic.003G040900) was exclusive to subpop 3 and predicted to cause a missense mutation (Figure 7B). *SbGI* was a high confidence predicted target of SbABF2. Looking at the variance in WUE between reference (Ref) and alternate (Alt) alleles, we found that lines with the Alt allele showed significant reductions in WUE (Figure 7B), and this was also evident in dynamic WUE patterns over time in response to drought (Figure 7C). This polymorphism showed significant associations based on both statistical tests (*p* < 0.05). To confirm that this moderate-effect SNP in *SbGI* was indeed associated with a difference in WUE and a not a false discovery due to the Tukey HSD test being confounded by differences in sample sizes between groups, we compared WUE for 100,000 random samples of 30 lines containing the reference allele and 100,000 random samples of 30 lines containing the high-effect SNP in *SbGI*. This permutation test showed that the differences between the groups were significant (mean *p* = 0.03670659). Furthermore, to test that the SNP specifically was significant and not just associated with subpop 3, we performed this permutation test on a set of 12 random subpop 3-specifc SNPs in the genome. None of these randomly selected SNPs had a *p* value < 0.10.

*SbGI* was recently shown to be involved in photoperiodic flowering time in sorghum (Abdul-Awal et al., 2020), which is a conserved function. In Arabidopsis, *GI* has also been implicated in drought escape through regulation of flowering time in a ABA-dependent manner (Riboni et al., 2016), and more recently was shown to modulate transcription of nine-cis-epoxycarotenoid dioxygenase 3 (NCED3), a rate-limiting enzyme in ABA biosynthesis, a mechanism by which diurnal stomatal movement could be modulated (Baek et al., 2020). Our analysis of chromatin accessibility in the *SbGI* promoter showed several upstream genomic regions that were regulated in response to drought and harbor ABF binding sites, as well as binding sites for several other stress-related TFs (Figure 7D). Candidate regulators of *SbGI* that potentially bind these sites were predicted based on interrogation of our drought-responsive GRN and were DE in response to drought, as were closely co-regulated genes in the SbGI sub-network (Figure 7E; Supplemental Data Set 14). Several of these close network first-neighbors to *SbGI* have putative roles in stomatal opening and dynamics.

We further tested whether variation in chromatin accessible regions were associated with certain sorghum subpops. Specifically, we looked at variation present in ABF binding sites within DARs across the genome and found clusters of SNPs that were specifically associated with one or more subpops (Supplemental Figure 10; Supplemental Data Set 15). Among these, several were found to potentially disrupt the core ABF binding sequence itself. Based on the SNP distribution across the subpops, we tested 115 SNPs for potential associations with WUE. Of these, five were found to be significantly associated with WUE (p < 0.05; Supplemental Data Set 15). Two of these associated with subpop 3 and their location within the core ABRE element could potentially cause disruption in ABF binding to the promoter region of a NAC TF (Sobic.005G064600) and a Light-harvesting chlorophyll B-binding protein (*Lhcb3/7*; Sobic.009G234600) (Supplemental Figure 10). The NAC TF was among high confidence predicted targets of SbABF1 and SbABF4 (Supplemental Data Set 11). *Lhcb* family members have also been implicated in stomatal response to ABA signaling (Xu et al., 2012; Liu et al., 2013). It is reasonable to expect that these polymorphisms could affect gene regulation and potentially ABA-mediated WUE.

## DISCUSSION

Global demand for cereals is expected to reach 3 billion tons in 2050, with increased demand largely coming from developing countries in Asia and Africa (Alexandratos and Bruinsma, 2012). In addition, yields have begun to plateau for major cereals crops (Ray et al., 2012; Grassini et al., 2013). To mitigate this, step changes are needed in crop improvement, and climate-resilient species are an untapped resource for novel genetic and genomic information (Reynolds et al., 2016; Varshney et al., 2018a). There have been some successes using biotechnology approaches to overexpress drought-responsive genes in crops to enhance drought tolerance (Hu and Xiong, 2014; Shi et al., 2015; Lamaoui et al., 2018). Often, however, these result in growth defects or yield losses, particularly in favorable environments. Accelerating crop yield to meet these growing global demands is going to require an understanding of the mechanisms governing existing traits that can be leveraged for enhancing productivity (Bailey-Serres et al., 2019). Our analysis of regulatory components across the sorghum genome and in response to a controlled drought stress provides a resource for interrogating the mechanisms of gene regulation in this stress resilient crop.

### Molecular features of ABA-response pathways and physiological mechanisms they control

Our analyses in sorghum highlighted the induction of several conserved drought-responsive pathways, including components of ABA synthesis, signaling and response. Many of the associated genes in these pathways were differentially regulated in response to drought, and we identified *cis*-elements that potentially drive their expression through TF-DNA interactions. In general, we observed more up-regulation in drought-responsive genes, which was consistent with enhanced chromatin accessibility. This suggests transcriptional activation genome-wide, which is the general mode of action in ABA-dependent signaling cascades. The ABA-dependent drought responses are propagated largely through the action of ABF/ABRE TFs, which are induced via phosphorylation by ABA-responsive SnRK2s. This phosphorylation allows for the rapid action of ABFs to transduce the drought response. The ABA-mediated ABF regulome has been largely mapped out in Arabidopsis (Song et al., 2016), and it is expected to be conserved to some degree across species. However, our specific knowledge of this is lacking in crops; for example, any diversification or specialization in regulatory function can be control points for modulating drought response.

Our network analyses revealed four ABF TFs (three positioned as hubs) in the drought-associated sub-network. A number of genes were predicted as direct targets of at least two ABFs, including several heat shock factors (HSFs), a G-box binding factor (GBF3), and notably, several core ABA signaling components such as 2C protein phosphatases (PPCs), including a homologs of *HAB1* and *HAI1* (Figure 6; Supplemental Data Set 11). There were also distinct sets of targets. Interestingly, SbAREB1/ABF2 was predicted to directly target genes related to flowering and the clock, including *SbGI* and the ortholog of *EARLY FLOWERING 3* (*ELF3;* Sobic.003G191700). ABA regulation of photoperiod genes including *GI* has been shown in Arabidopsis (Riboni et al., 2016). Photoperiodicity is important in sorghum breeding, and underlies differences between grain and bioenergy types. Our analysis of the *SbGI* locus based on pan-genomic data revealed specific variation that was significantly associated with decreased WUE. Recently, a study in Arabidopsis showed that GI interacted with the TF, ENHANCED EM LEVEL (EEL)/basic Leu zipper 12, to regulate diurnal expression of NCED3, a rate-limiting enzyme in ABA biosynthesis (Baek et al., 2020). This regulation was proposed to enhance drought tolerance through regulation of ABA production and diurnal control of stomatal opening to reduce transpiration during the day. Our analysis of the SbGI subnetwork showed several strong network connections as clear links to ABA-mediated drought response including orthologs of the PPC *HAB1* and drought tolerance gene *XERICO* (Ko et al., 2006), as well as genes encoding osmoregulatory LEA proteins. In addition, several genes that were strongly co-regulated with *SbGI* have been implicated in stomatal dynamics; for example, *ARABIDOPSIS PROTEIN KINASE 1b* (*APK1b*) is predominantly expressed in guard cells and is required for full light-induced stomatal opening in Arabidopsis (Elhaddad et al., 2014), and *ACTIN DEPOLYMERIZING FACTORs* (*ADFs*) have been implicated in guard cell movement and regulated by ABA through ABFs (Qian et al., 2019; Yang et al., 2020).

Our finding that allelic variation in *SbGI* contributes to lower WUE suggests that this genetic variation, or variation in the regulation of *SbGI*, can be leveraged for improving crop tolerance to drought stress. The apparent specificity for SbABF2 in the regulation of *SbGI* provides insights into how pathways such as flowering time are intimately linked through molecular interactions in the drought response network. Unraveling the spatiotemporal action of these drought-responsive TFs, e.g., the ABFs, and their shared and unique predicted targets, provide clues on how to regulate certain aspects of drought response in sorghum independently of other connections in the overall network in which they reside.

### Regulatory links between ABA and nutrient pathways

ABA is an integral player in the physiological response to drought since it mitigates water loss through stomatal conductance, increases root surface area, and induces the production of osmoprotectants. Therefore, understanding how ABA-dependent and independent regulation interface at the molecular level with these processes is critical to enhancing water use and photosynthetic efficiencies in crops. There has been extensive characterization in Arabidopsis of overlaps in ABA and sugar sensing and signaling pathways, and some insights from other systems (León and Sheen, 2003; Eveland and Jackson, 2012). Sugar sensing pathways signal spatiotemporal carbohydrate status and feedback regulate photosynthesis, sugar transport, assimilation and storage. Our functional enrichment analyses revealed overrepresentation of differentially regulated genes associated with responses to both sucrose and to ABA, and were positively regulated in both leaf and root tissues. We also observed dynamic expression patterns of genes related to ABA and sucrose metabolism across the developmental gradient of the leaf in response to drought. For example, a gene encoding *Zeaxanthin epoxidase* (*zep*; Sobic.006G097500), which functions in the first step of ABA biosynthesis, is up-regualed in response to drought and increases in expression from base to tip of the leaf, while *sucrose synthase* (*sus*; Sobic.001G344500), involved in reversible sucrose metabolism, is down-regulated in response to drought and shows an expression trajectory directly opposite that of *zep* across the leaf gradient (Figure 3E).

Functional enrichment analyses also indicated shifts in C and N balance in response to drought, which is expected given the physiological changes that accompany the response. Insights into the regulation of these pathways through specific TF-DNA interaction can be important targets for enhancing productivity. Signatures of altered C and N metabolism, transport and assimilation can provide insights into drought-induced leaf senescence, which occurs downstream of our analyses (Figure 1A; (Chen et al., 2015)). Interestingly, genes related to nitrate transport showed differential responses between leaf and root. We observed down-regulation of several transporters in the root in response to drought, including nitrate transporter *NRT2.5* (Sobic.003G188200). Alternatively, *asparagine synthase* (*asn;* Sobic.001G406800) was up-regulated across tissues, which would suggest increased N assimilation relative to uptake. The processes regulating water and N uptake and use must be tightly interconnected, yet currently largely not understood (Araus et al., 2020)

### Transcriptional networks of hormone cross-talk and environmental stimuli

In addition to ABA-related processes, there were signatures of regulation by BRs and ethylene, which have also been implicated in various aspects of abiotic stress response. Notably, genes associated with BR synthesis and response pathways were down-regulated in response to drought and also associated with less accessibility in open chromatin regions. It is known that there are a number of ways that ABA and BR pathways are antagonistic (Nolan et al., 2020). The negative regulator of BR signaling, BIN2, phosphorylates SnRK2s to activate ABA pathway genes and TFs like ABI5 (Cai et al., 2014; Hu and Yu, 2014). ABI1 and ABI2 dephosphorylate BIN2, but are repressed by ABA (Wang et al., 2018). They also interact through downstream transcription factors BES1 or ABI3 and ABI5. BIN2 has also been shown to regulate stomatal development antagonistically of ABA (Kim et al., 2012). Regulation of the BR pathway in sorghum is significant since it underlies at least one of the dwarfing genes used in sorghum breeding, whereas breeding programs in other commercial cereal crops largely leverage the GA pathway (Ordonio et al., 2014; Hirano et al., 2017).

Our study was performed in a controlled growth environment, and we could isolate the drought response more accurately by controlling for other variables. Of course, in a drought-stressed field environment, the molecular and physiological responses become more complex through interactions with other stresses that are typically coupled with drought, e.g., heat, salt or low nutrient stresses. Molecular signaling components of these stress pathways can also overlap, and perhaps rewiring drought-specific responses to some degree. Our functional enrichment analyses showed that DE genes in response to drought were associated with several types of stress and stress hormones, suggesting reuse of many genes in various stress contexts. Several of these are TFs that interface stress response pathways such as cold, heat, and salt (Nakashima et al., 2014). This study provides a set of drought-responsive promoters and predicted TF-DNA interactions that can be leveraged in manipulating a plant’s response to its environment.

Our results provide a first look at the drought responsive genome in stress resilient sorghum. We observed both known and novel aspects of drought response and inferred regulatory modules that function at the core of this response in young sorghum plants. Sorghum boasts a wide range of adaptation to various local climates, which likely underlies its stress tolerance and provides a repository for identifying novel response loci. Extending these analyses to other genotypes will further enhance our understanding of how natural variation regulates response to the environment at the molecular level. Leveraging extensive genetic diversity in sorghum can help pinpoint functional variants in gene regulation, e.g., regulatory elements that improve stress resilience with minimal disruption to the complex gene networks governing plant growth and development. Elucidating the regulatory components that control stress response in resilient crops such as sorghum will help identify ways to fine-tune these pathways and improve stress resilience in more susceptible crop species.

## METHODS

### Experimental design and tissue collection

BTx623 sorghum seeds were germinated on peat moss in petri dishes for 3 days and then transplanted into a Metromix360/Turface MVP blend soil in 9×2×2 tree pots. Plants were grown in a controlled, high-light chamber at the Danforth Center Integrated Plant Growth Facility in 500μMol light, 14 hour days, 28°C days, 24°C nights. Pots were weighed each morning and drought stressed plants were watered based on what they lost the previous day to transpiration. Soil moisture content (SMC) and field capacity was measured by drying soil in a 37°C drying chamber for 4 days, weighing out material and adding water to retention capacity and re-weighing. Total water weight was calculated, and used as reference for appropriate watering targets. From 12-14 DAS, 10 mL of the respective solution was applied and pots were maintained at 80% SMC. From 15-23 DAS, plants were watered with reverse osmosis (RO) water and maintained at either 80% SMC (WW) or 25% (WS). Plants were sampled at 23 DAS in a random order, 1 hour after lights on and ~22 hrs since the previous watering. The emerging leaf was carefully dissected down to the ligular region and the developing leaves, the leaves within the emerging leaf, were removed and segmented into six 1 cm sections (1,2,3 cm above the ligule; 2 cm above and 2 cm below the midpoint or where the leaf had emerged from the whorl; and 4 cm from the tip of the leaf) were dissected from the plants and flash frozen in liquid N. The base of the stem was pulled from soil to remove crown roots. Roots were quickly washed, dried, and flash frozen in liquid N.

### RNA-seq library preparation

Total RNA was extracted from the ~100 mg of ground tissue (inner leaf and root) using an in-house Trizol extraction protocol. DNA contamination was removed using the turbo DNA-free kit (invitrogen) following the manufacturers settings and the RNA quality was checked using the Agilent Bioanalyzer RNA chip. RNA libraries were prepared using ~950 ng of the DNAase treated total RNA using the NEBNext Ultra Directional RNA Library Prep Kit (Illumina) and size selected for 200bp insert size. Adapters supplied in the NEBNext kit were used and 12 PCR cycles were used for the cDNA amplification. The final libraries were quality checked on an Agilent bioanalyzer using a DNA 1000 chip and 100bp SE reads were generated using the Illumina HiSeq 4000 platform (University of Illinois at Urbana-Champaign W.M. Keck Center). We obtained an average of 45 million reads per library.

For the leaf gradient samples, we generated 32 RNA-seq libraries using the Lexogen Quantseq 3’ mRNA-seq Library Prep Kit FWD for Illumina (cat # 015.384) following the manufacturer’s protocol and amplified using 12 cycles. Libraries were sequenced to an average 4M reads per library.

### Analysis of RNA-seq data

Single-end (SE) 100 nt reads (and paired end 150 nt) were adapter- and quality-trimmed using Trim Galore with parameters:--stringency 3 (--paired) --clip_R1 13 (--clip_R2 13) --length 20 -q 20. The *S. bicolor* genome sequence (v3.0.1) and annotation v3.1.1 were obtained from Phytozome. Transcript quantification was performed using the quasi-mapping method, Salmon (parameters: **SE** --incompatPrio 0.0 --seqBias --libType SR; **PE** --incompatPrio 0.0 --allowDovetail --seqBias --gcBias --libType A). Differential expression (DE) analysis was performed using the edgeR package in R (parameters: glmfit, glmLRT, robust = T). An FDR cutoff of 0.05 and at least two fold change in expression between treatments was considered significantly DE. To visualize on the JBrowse genome browser, reads were mapped to the genome using the STAR package (parameters: --sjdbOverhang 100, --twopassMode Basic, --outFilterMultimapNmax 10, --alignSJoverhangMin 8, --alignSJDBoverhangMin 1, --outFilterMismatchNmax 999, --outFilterMismatchNoverReadLmax 0.04, --alignIntronMin 20 --alignIntronMax 1000000, --alignMatesGapMax 1000000). BAM files were converted to bigwig using bamCoverge (deepTools) with parameters --binSize 1 --normalizeUsing RPKM prior to loading on the genome browser.

RNAseq from leaf gradient samples were performed using Quantseq 3’ FWD kit. As read2 of the read pairs primarily contains PolyA sequences, only read1 was used for further processing. Reads were adapter- and quality-trimmed using Trim Galore with parameters:--stringency 3 --clip_R1 13 --length 20 -q 20 followed by a second run with --stringency 3 -a A{10} --length 20 -q 20 in order to remove any PolyA sequences. Quantseq 3’ end sequencing method typically represents reads near the 3’ end of transcripts. We observed several instances of poorly annotated genes with completely missing or a short 3’ UTR which led to reads mapping out of the genes. To minimize such issues, we curated the gene models by extending the 3’ end by 250 nt or 750 nt for the genes with or without annotated 3’ UTR respectively. Transcript quantification was performed using the quasi-mapping method, Salmon (parameters: --validateMappings --incompatPrio 0.0 --noLengthCorrection --noEffectiveLengthCorrection --noFragLengthDist --libType SF).

### Nuclei isolation and ATAC-seq library construction

Flash-frozen inner leaf and crown root tissue were ground in liquid N and ~0.2-0.3 g of tissue was aliquoted into a 15 mL falcon tube. Ground tissue was resuspended in 4 mL of 1x nuclei isolation buffer (16 mM HEPES; pH8, 200 mM sucrose, 0.8 mM MgCl2, 4 mM KCl, 32 % Glycerol, 0.25% Triton X-100, 1x complete protease inhibitor, 0.1% 2-ME, 0.1 mM PMSF) very gently at 4°C for 20 minutes and then filtered through 2 sheets of mira cloth. The resulting elutent (~3 mL) was equally split into two 2 mL eppendorf tubes and centrifuged at 1,000 x g for 15 minutes. After discarding the supernatant, the nuclei pellets in the two eppendorf tubes were resuspended in 400 μL of 1x tagmentation buffer, combined into a single tube and centrifuged at 1,000 x g for 5 minutes as a wash step (total 2 x washes). The nuclei were resuspended in 100 μL of 1x tagmentation buffer and observed under a microscope (2% acetocarmine stain) to check nuclear integrity. Nuclei were counted using a hemocytometer, and approximately 50,000 nuclei were used for tagmentation.

Tagmentation was performed using the Illumina DNA Library Prep Kit (FC-121-1031) and Index Kit (FC-121-1011) with 2.5 μL Tn5 enzyme, 2.5 μL of 2x tagmentation buffer (20 mM Tris Base, 10 mM MgCl2, 20% v/v dimethylformamide), and 20 μL of nuclei for each sample and incubated for 1 hr at 37°C. To each reaction, added 22 μL of H2O, 2.5 μL 10% SDS, and 0.5 μL of Proteinase K and incubated at 55 °C for 1 hr. Tagmented libraries were purified using the Zymo clean and concentrator kit, eluting with RSB buffer. DNA was quantified using Qubit. 25 ng of tagmented DNA was combined with 2.5 μL of both index primers, 7.5 μL PCR master mix, 2.5 μL NPM and filled to 25 μL with RSB buffer. Libraries were amplified with 12 cycles. Libraries were diluted to 50 μL and cleaned up with a two-sided Ampure XP bead size selection. A 0.5:1 bead:sample ratio followed by a 1.2:1 bead:sample ratio was used to select ~200-1000 bp libraries. Sequencing was performed using Illumina HiSeq 4000 paired-end (PE) 50 platform.

### ATAC-seq mapping and peak calling

Paired-end 50 bp reads were trimmed to remove adapters using BBDuk (qtrim=r, trimq=6, minlen=1; https://jgi.doe.gov/data-and-tools/bbtools/). Adapter-trimmed reads were mapped to sorghum chromosome (v.3.0.1 from Phytozome (https://phytozome.jgi.doe.gov/), sorghum mitochondria (NCBI reference: NC_008360.1) and sorghum chloroplast (NCBI reference: NC_008602.1)) genomes using *bowtie2 (Langmead and Salzberg, 2012)* with settings `--sensitive --maxins 2000’. Reads were filtered to remove those below MAPQ < 20, multi-mapping reads and those that mapped to chloroplast and mitochondria genomes. Duplicate reads were filtered using Picard (https://broadinstitute.github.io/picard/) with the settings: ‘MarkDuplicates MAX_FILE_HANDLES_FOR_READ_ENDS_MAP=1000 PG=null MAX_RECORDS_IN_RAM=5000000 REMOVE_DUPLICATES=true’. To visualize in Jbrowse, BAM files were converted to bigwig using deeptools bamCoverage (Ramírez et al., 2014).

ATAC peaks were called using Hypergeometric Optimization of Motif EnRichment (HOMER) (Heinz et al., 2010) with settings ‘-gsize 7.09e8 -region -size 150 -tbp 0 -localSize 5000 -L 4 -fdr 0.01’. Peaks present in at least two biological replicates (minimum 25% overlap) were determined as high-confidence peaks. High-confidence peaks that overlapped by at least 25% between treatments (WW and WS) were merged using *bedops*. High-confidence peaks that did not overlap by a minimum of 25% were considered unique to the respective condition. Both of these peak sets constitute ‘union peaks’ used for determining differential accessibility (DARs). Read counts (transposase insertion sites) within union peaks were calculated using custom scripts. The spearman correlation among replicates for all high-confidence ATAC regions in inner leaf and crown root was calculated using the plotCorrelation function in *deeptools* (Ramírez et al., 2014) on the coverage (bigwig) files. Our sequence depth was higher for inner leaf tissue, so we called more unique peaks overall compared to the crown roots. DARs were calculated using EdgeR (settings: glmfit, robust = FALSE, glmLRT (), FDR < 0.05).

The distribution of the high confidence ATAC-seq peaks and DARs across gene features was performed using custom scripts using R package *Genomic Ranges*. The midpoint of the ATAC-seq peaks were used for assignment to each of the gene features. Primary gene transcripts of the *S. bicolor* v3.1.1 annotations were used. The first gene within a 10kb distance (on either side of DAR) was considered to be associated with that DAR.

### GO enrichment analyses

Gene Ontology enrichment analysis was performed by Clusterprofiler package (v 3.10.1) in R (Yu et al., 2012). A custom GO annotation generated by GOMAP pipeline was used. Significantly enriched GO categories were identified with the following parameter of enricher function: pvalueCutoff = 0.05, pAdjustMethod = “BH”, minGSSize = 10, maxGSSize = 1000, qvalueCutoff = 0.05.

### Motif analysis

*De novo* motif analysis was performed using the Discriminative Regular Expression Motif Elicitation (DREME) tool in the MEME suite (Bailey, 2011) with default settings. Random genomic fragments of equal size to DARs from all ATAC HS regions combined were used as the background control. Resulting *de novo* motifs were compared to the JASPAR plant database (2018) to identify the closest matches using TOMTOM (Gupta et al., 2007). To compare *de novo* motifs across tissues and treatment groups, we used the STAMP (Mahony and Benos, 2007) web interface (http://www.benoslab.pitt.edu/stamp/). Genomic locations of *de novo* motifs were identified using Find Individual Motif Occurrences (Bailey et al., 2009) with settings “ --max-strand --parse-genomic-coord --max-stored-scores 1000000 --thresh 0.0005”. The background file for FIMO was generated using all the accessible peaks.

### Gene co-expression network analysis and target prediction

A gene co-expression network was constructed using the WGCNA v1.69 package in R (Langfelder and Horvath, 2008). Expression data (TPM) from inner leaf, leaf gradient and root (44 samples) was used as input. Genes with low expression (< 5 TPM in at least 3 samples) were filtered out. Expression values from 22,847 genes were log2 transformed and a signed network was created using following parameters: corType=“pearson”, maxBlockSize=25000, networkType=“signed”, power=14, minModuleSize=30, deepSplit=2, mergeCutHeight=0.25). Based on the scale free topology model fit, a softthresholding power of 14 was selected. The network identified 23 modules of genes with distinct eigengenes each represented by a different color. The relationship between modules (module eigengene) and traits was determined and reflected as correlation coefficient and p-value. Treatments (WW and WS) as well as tissue types (inner leaf, base+2, m+2, p+4, tip and root) were considered as traits. Module-trait correlations of R^2^ > 0.5 were considered as highly correlated. Hub genes for each module were identified by selecting genes with module membership > 0.8.

Putative target genes of TFs in the drought modules were identified by R implementation of GENIE3 (Huynh-Thu et al., 2010) using the expression values with following parameters: treeMethod= “RF”, K= “sqrt”, nTrees= 1000. Regulatory links with weight > 0.005 were used for further analysis. Target genes were filtered by those that were DAR proximal and DE followed by the presence of corresponding TF binding motifs (obtained from the DREME and scanned by FIMO described above) in the associated DARs. Target network models for SbAREB1/ABF2 and SbABF3 were created using select filtered targets in the igraph R package (Csardi et al., 2006).

### Controlled environment phenotyping of a sorghum diversity panel in response to drought

The sorghum diversity panel consisted of 380 genotypes from the BAP. Three plants per genotype were subjected to two watering regimes (100% field capacity for WW and 30% field capacity for WS). Plants were sown in a soil mixture (ProMix BRK20 + 14-14-14 Osmocote (1.5 lb/cu yd)) and acclimatized in a controlled growth chamber for 10 days before being transferred to the LemnaTec Scanalyzer phenotyping system at the Danforth Center Bellwether Foundation Phenotyping Facility. All plants were maintained at 100% field capacity for two days and at 12 days after planting (DAP) the WS watering regime was initiated for half of the plants. Automated weighing and watering two times daily maintained plants to desired field capacity. Visible spectrum (VIS) images were taken each day (2 side views). PlantCV and custom Python scripts were used to color-correct and analyze the VIS images and to calculate WUE in both WW and WS conditions. To accommodate the total set of 380 lines, two runs of 190 lines were carried out back to back.

### Sorghum population structure and pan-genome analyses

The TASSEL 5 pipeline (Glaubitz et al., 2014) was used to generate a hapmap file based on GBS data from the BAP. Plink (Rentería et al., 2013) was used to create a binary biallelic genotype table (.bed file) which was then used in the ADMIXTURE program (Alexander et al., 2009). ADMIXTURE was run in a loop, in which the population (K) value was modified from 3 to 10 and a K of 8 was chosen based on the Cross Validation Error.

The PAV and SNP effect files were generated based on whole genome resequencing data from the sorghum BAP (https://terraref.org). Whole genome shotgun sequencing data from 363 genotypes was used for the SNP effect analyses. Of the 363 sequenced genotypes, only 205 genomes had a depth of coverage sufficient for reliable PAV prediction. The differences in PAV and SNPs between the drought resilient and susceptible genotypes within genes and in DARs were identified using custom R scripts.

## DATA ACCESSIBILITY

Raw data have been deposited in NCBI SRA as Bioproject ID PRJNA686818 and will be available upon publication. Processed data and metadata are available at NCBI GEO accession # GSE163578 and will be released upon publication.

## AUTHOR CONTRIBUTIONS

ALE designed and advised the research. MB performed the ATAC-seq and RNA-seq experiments. RKP, IK and PO performed data analyses. TCM led and advised phenotyping and pan-genome analyses. RKP, IK and ALE wrote the paper. All authors reviewed and edited the final manuscript.

## ACKNOWLEDGEMENTS

The authors would like to thank Edoardo Bertolini in the Eveland lab for input on the drought experiment and network analyses, Sarit Weissmann in the Mockler lab for advice on nuclei isolation, and Kerry Bubb in Dr. Christine Queitsch’s lab at the University of Washington for advice on quantification on differentially accessible chromatin. Thank you to Kevin Reilly and his team at the DDPSC Integrated Plant Growth Facility for chamber maintenance and plant care, Noah Fahlgren and his team in the Data Science core at DDPSC for cluster maintenance, and to Mindy Darnell and Leo Chavitz in the DDPSC Phenotyping core. We also thank Scott Lee (Mockler lab) and Adam Healey and Jeremy Schmutz (HudsonAlpha) for sorghum BAP resequencing and pan-genomic analyses, which was funded by DOE-JGI and ARPA-E award #DE-AR0000594 to TCM. The rest of the work presented here was funded by the U.S. Department of Energy BER awards #DE-SC0014395 and #DE-SC0020401 to ALE and TCM.

## SUPPLEMENTAL MATERIAL

### Supplemental Figures

**Supplemental Figure 1.** Nuclei isolation, nucleosome phasing, and ATAC-seq library QC.

**Supplemental Figure 2.** RNA-seq data showed strong correlations among biological replicates and separation of tissue types by PCA.

**Supplemental Figure 3.** Up-regulated genes showed enhanced chromatin accessibility around their proximal promoters in response to drought stress.

**Supplemental Figure 4.** WGCNA module identification and relationship.

**Supplemental Figure 5.** WGCNA module eigengene (ME) expression plots for all modules.

**Supplemental Figure 6.** Plots showing cumulative water usage across the BAP in WW and WS controlled environment conditions.

**Supplemental Figure 7.** Heatmap of WUE residuals over the drought response period for genotypes in subpops 2 and 3 from controlled environment phenotyping experiment.

**Supplemental Figure 8.** Presence/Absence Variation (PAV) in ABF target genes.

**Supplemental Figure 9.** SNP variation within ABF target genes.

**Supplemental Figure 10.** SNP variation within ABRE motifs in DARs.

### Supplemental Tables

**Supplemental Table 1.** Mapping statistics for ATAC-seq data from inner leaf and crown root.

**Supplemental Table 2.** Homer peak call statistics based on ATAC-seq data for the inner leaf and crown root samples.

**Supplemental Table 3.** Distribution of high confidence ATAC-seq peaks across genomic features.

### Supplemental Data Sets

**Supplemental Data Set 1.** File with all the HOMER peaks per replicates and combined.

**Supplemental Data Set 2**. All the high confidence HOMER peaks are listed.

**Supplemental Data Set 3.** Location of DARs (between control and drought; leaf and root), distance and location to closest gene(s).

**Supplemental Data Set 4.** Supp xls file with all genes and expression values/statistics for comparison.

**Supplemental Data Set 5.** GO enrichment for DE genes proximal to DARs (root and leaf) - quadrants.

**Supplemental Data Set 6.** Leaf gradient RNAseq data (TPM).

**Supplemental Data Set 7.** PWM enriched in the DARs sub-divided by positive and anti-correlation to the DE gene (within 10 kb distance).

**Supplemental Data Set 8.** List of DE TFs and their expression values whose PWM signatures are enriched in the de novo motif analysis.

**Supplemental Data Set 9.** File with drought module members along with gene annotations, proximal DARs and GO enrichment, WGCNA connections and GENIE3 predicted targets of TFs in the drought modules.

**Supplemental Data Set 10.** GO enrichment of ABF2 and ABF3 predicted target genes.

**Supplemental Data Set 11.** Lists of high confidence target genes of all 4 ABF TFs.

**Supplemental Data Set 12.** WUE over time under 2 conditions for all genotypes.

**Supplemental Data Set 13.** PAV and SNP variation in ABF target genes.

**Supplemental Data Set 14.** First neighbor connections of GI in the co-expression network and GO enrichment analyses.

**Supplemental Data Set 15.** SNPs inside predicted ABRE motifs across all DARs.

## REFERENCES

Abdul-Awal, S.M., Chen, J., Xin, Z., and Harmon, F.G. (2020). A sorghum gigantea mutant attenuates florigen gene expression and delays flowering time. Plant Direct 4: e00281.

Alexander, D.H., Novembre, J., and Lange, K. (2009). Fast model-based estimation of ancestry in unrelated individuals. Genome Res. 19: 1655–1664.

Alexandratos, N. and Bruinsma, J. (2012). World agriculture towards 2030/2050: the 2012 revision. ESA Working paper No. 12-03. Rome, FAO.

Amara, I., Capellades, M., Ludevid, M.D., Pagès, M., and Goday, A. (2013). Enhanced water stress tolerance of transgenic maize plants over-expressing LEA Rab28 gene. J. Plant Physiol. 170: 864–873.

Araus, V., Swift, J., Alvarez, J.M., Henry, A., and Coruzzi, G.M. (2020). A balancing act: how plants integrate nitrogen and water signals. Journal of Experimental Botany 71: 4442–4451.

Baek, D. et al. (2020). The GIGANTEA-ENHANCED EM LEVEL Complex Enhances Drought Tolerance via Regulation of Abscisic Acid Synthesis. Plant Physiol. 184: 443–458.

Bailey-Serres, J., Parker, J.E., Ainsworth, E.A., Oldroyd, G.E.D., and Schroeder, J.I. (2019). Genetic strategies for improving crop yields. Nature 575: 109–118.

Bailey, T.L. (2011). DREME: motif discovery in transcription factor ChIP-seq data. Bioinformatics 27: 1653–1659.

Bailey, T.L., Boden, M., Buske, F.A., Frith, M., Grant, C.E., Clementi, L., Ren, J., Li, W.W., and Noble, W.S. (2009). MEME SUITE: tools for motif discovery and searching. Nucleic Acids Res. 37: W202–8.

Battaglia, M., Olvera-Carrillo, Y., Garciarrubio, A., Campos, F., and Covarrubias, A.A. (2008). The enigmatic LEA proteins and other hydrophilins. Plant Physiol. 148: 6–24.

Benlloch, R. and Lois, L.M. (2018). Sumoylation in plants: mechanistic insights and its role in drought stress. J. Exp. Bot. 69: 4539–4554.

Boyles, R.E., Brenton, Z.W., and Kresovich, S. (2019). Genetic and genomic resources of sorghum to connect genotype with phenotype in contrasting environments. Plant J. 97: 19–39.

Brenton, Z.W., Cooper, E.A., Myers, M.T., Boyles, R.E., Shakoor, N., Zielinski, K.J., Rauh, B.L., Bridges, W.C., Morris, G.P., and Kresovich, S. (2016). A Genomic Resource for the Development, Improvement, and Exploitation of Sorghum for Bioenergy. Genetics 204: 21–33.

Brkljacic, J. and Grotewold, E. (2017). Combinatorial control of plant gene expression. Biochim. Biophys. Acta Gene Regul. Mech. 1860: 31–40.

Buenrostro, J.D., Wu, B., Chang, H.Y., and Greenleaf, W.J. (2015). ATAC-seq: A Method for Assaying Chromatin Accessibility Genome-Wide. Curr. Protoc. Mol. Biol. 109: 21.29.1–9.

Cai, Z., Liu, J., Wang, H., Yang, C., Chen, Y., Li, Y., Pan, S., Dong, R., Tang, G., Barajas-Lopez, J. de D., Fujii, H., and Wang, X. (2014). GSK3-like kinases positively modulate abscisic acid signaling through phosphorylating subgroup III SnRK2s in Arabidopsis. Proc. Natl. Acad. Sci. U. S. A. 111: 9651–9656.

Chandra Babu, R., Zhang, J., Blum, A., David Ho, T.-H., Wu, R., and Nguyen, H.T. (2004). HVA1, a LEA gene from barley confers dehydration tolerance in transgenic rice (Oryza sativa L.) via cell membrane protection. Plant Sci. 166: 855–862.

Chen, D., Wang, S., Xiong, B., Cao, B., and Deng, X. (2015). Carbon/Nitrogen Imbalance Associated with Drought-Induced Leaf Senescence in Sorghum bicolor. PLoS One 10: e0137026.

Christmann, A., Grill, E., and Huang, J. (2013). Hydraulic signals in long-distance signaling. Current Opinion in Plant Biology 16: 293–300.

Christmann, A., Weiler, E.W., Steudle, E., and Grill, E. (2007). A hydraulic signal in root-to-shoot signalling of water shortage. Plant J. 52: 167–174.

Csardi, G., Nepusz, T., and Others (2006). The igraph software package for complex network research. InterJournal, complex systems 1695: 1–9.

Duan, J. and Cai, W. (2012). OsLEA3-2, an abiotic stress induced gene of rice plays a key role in salt and drought tolerance. PLoS One 7: e45117.

Elhaddad, N.S., Hunt, L., Sloan, J., and Gray, J.E. (2014). Light-induced stomatal opening is affected by the guard cell protein kinase APK1b. PLoS One 9: e97161.

Endo, A. et al. (2008). Drought induction of Arabidopsis 9-cis-epoxycarotenoid dioxygenase occurs in vascular parenchyma cells. Plant Physiol. 147: 1984–1993.

Eveland, A.L. and Jackson, D.P. (2012). Sugars, signalling, and plant development. J. Exp. Bot. 63: 3367–3377.

Fahlgren, N. et al. (2015). A Versatile Phenotyping System and Analytics Platform Reveals Diverse Temporal Responses to Water Availability in Setaria. Mol. Plant 8: 1520–1535.

Fujita, Y., Yoshida, T., and Yamaguchi-Shinozaki, K. (2013). Pivotal role of the AREB/ABF-SnRK2 pathway in ABRE-mediated transcription in response to osmotic stress in plants. Physiologia Plantarum 147: 15–27.

Glaubitz, J.C., Casstevens, T.M., Lu, F., Harriman, J., Elshire, R.J., Sun, Q., and Buckler, E.S. (2014). TASSEL-GBS: a high capacity genotyping by sequencing analysis pipeline. PLoS One 9: e90346.

Grassini, P., Eskridge, K.M., and Cassman, K.G. (2013). Distinguishing between yield advances and yield plateaus in historical crop production trends. Nat. Commun. 4: 2918.

Gupta, A., Rico-Medina, A., and Caño-Delgado, A.I. (2020). The physiology of plant responses to drought. Science 368: 266–269.

Gupta, S., Stamatoyannopoulos, J.A., Bailey, T.L., and Noble, W.S. (2007). Quantifying similarity between motifs. Genome Biol. 8: R24.

Han, J., Wang, P., Wang, Q., Lin, Q., Chen, Z., Yu, G., Miao, C., Dao, Y., Wu, R., Schnable, J., Tang, H., and Wang, K. (2020). Genome-wide Characterization of DNase I-hypersensitive Sites and Cold Response Regulatory Landscapes in Grasses. Plant Cell.

Heinz, S., Benner, C., Spann, N., Bertolino, E., Lin, Y.C., Laslo, P., Cheng, J.X., Murre, C., Singh, H., and Glass, C.K. (2010). Simple combinations of lineage-determining transcription factors prime cis-regulatory elements required for macrophage and B cell identities. Mol. Cell 38: 576–589.

Hirano, K., Kawamura, M., Araki-Nakamura, S., Fujimoto, H., Ohmae-Shinohara, K., Yamaguchi, M., Fujii, A., Sasaki, H., Kasuga, S., and Sazuka, T. (2017). Sorghum DW1 positively regulates brassinosteroid signaling by inhibiting the nuclear localization of BRASSINOSTEROID INSENSITIVE 2. Sci. Rep. 7: 126.

Hobo, T., Kowyama, Y., and Hattori, T. (1999). A bZIP factor, TRAB1, interacts with VP1 and mediates abscisic acid-induced transcription. Proc. Natl. Acad. Sci. U. S. A. 96: 15348–15353.

Hu, H. and Xiong, L. (2014). Genetic engineering and breeding of drought-resistant crops. Annu. Rev. Plant Biol. 65: 715–741.

Huynh-Thu, V.A., Irrthum, A., Wehenkel, L., and Geurts, P. (2010). Inferring regulatory networks from expression data using tree-based methods. PLoS One 5(9): e12776

Hu, Y. and Yu, D. (2014). BRASSINOSTEROID INSENSITIVE2 interacts with ABSCISIC ACID INSENSITIVE5 to mediate the antagonism of brassinosteroids to abscisic acid during seed germination in Arabidopsis. Plant Cell 26: 4394–4408.

Jin, J., Tian, F., Yang, D.-C., Meng, Y.-Q., Kong, L., Luo, J., and Gao, G. (2017). PlantTFDB 4.0: toward a central hub for transcription factors and regulatory interactions in plants. Nucleic Acids Res. 45: D1040–D1045.

Joshi, R., Wani, S.H., Singh, B., Bohra, A., Dar, Z.A., Lone, A.A., Pareek, A., and Singla-Pareek, S.L. (2016). Transcription Factors and Plants Response to Drought Stress: Current Understanding and Future Directions. Front. Plant Sci. 7: 1029.

Kim, T.-W., Michniewicz, M., Bergmann, D.C., and Wang, Z.-Y. (2012). Brassinosteroid regulates stomatal development by GSK3-mediated inhibition of a MAPK pathway. Nature 482: 419–422.

Ko, J.-H., Yang, S.H., and Han, K.-H. (2006). Upregulation of an Arabidopsis RING-H2 gene, XERICO, confers drought tolerance through increased abscisic acid biosynthesis. Plant J. 47: 343–355.

Lamaoui, M., Jemo, M., Datla, R., and Bekkaoui, F. (2018). Heat and Drought Stresses in Crops and Approaches for Their Mitigation. Frontiers in Chemistry 6.

Langfelder, P. and Horvath, S. (2008). WGCNA: an R package for weighted correlation network analysis. BMC Bioinformatics 9: 559.

Langmead, B. and Salzberg, S.L. (2012). Fast gapped-read alignment with Bowtie 2. Nat. Methods 9: 357–359.

Lasky, J.R. et al. (2015). Genome-environment associations in sorghum landraces predict adaptive traits. Sci Adv 1: e1400218.

León, P. and Sheen, J. (2003). Sugar and hormone connections. Trends Plant Sci. 8: 110–116.

Li, P., Ponnala, L., Gandotra, N., Wang, L., Si, Y., Tausta, S.L., Kebrom, T.H., Provart, N., Patel, R., Myers, C.R., and Others (2010). The developmental dynamics of the maize leaf transcriptome. Nat. Genet. 42: 1060.

Liu, R., Xu, Y.-H., Jiang, S.-C., Lu, K., Lu, Y.-F., Feng, X.-J., Wu, Z., Liang, S., Yu, Y.-T., Wang, X.-F., and Zhang, D.-P. (2013). Light-harvesting chlorophyll a/b-binding proteins, positively involved in abscisic acid signalling, require a transcription repressor, WRKY40, to balance their function. J. Exp. Bot. 64: 5443–5456.

Lu, Z., Marand, A.P., Ricci, W.A., Ethridge, C.L., Zhang, X., and Schmitz, R.J. (2019). The prevalence, evolution and chromatin signatures of plant regulatory elements. Nat Plants.

Maher, K.A. et al. (2018). Profiling of Accessible Chromatin Regions across Multiple Plant Species and Cell Types Reveals Common Gene Regulatory Principles and New Control Modules. Plant Cell 30: 15–36.

Mahony, S. and Benos, P.V. (2007). STAMP: a web tool for exploring DNA-binding motif similarities. Nucleic Acids Res. 35: W253–8.

Mittal, S., Banduni, P., Mallikarjuna, M.G., Rao, A.R., Jain, P.A., Dash, P.K., and Thirunavukkarasu, N. (2018). Structural, Functional, and Evolutionary Characterization of Major Drought Transcription Factors Families in Maize. Front Chem 6: 177.

Morris, G.P. et al. (2013). Population genomic and genome-wide association studies of agroclimatic traits in sorghum. Proc. Natl. Acad. Sci. U. S. A. 110: 453–458.

Mullet, J., Morishige, D., McCormick, R., Truong, S., Hilley, J., McKinley, B., Anderson, R., Olson, S.N., and Rooney, W. (2014). Energy Sorghum—a genetic model for the design of C4 grass bioenergy crops. J. Exp. Bot. 65: 3479–3489.

Munemasa, S., Hauser, F., Park, J., Waadt, R., Brandt, B., and Schroeder, J.I. (2015). Mechanisms of abscisic acid-mediated control of stomatal aperture. Curr. Opin. Plant Biol. 28: 154–162.

Nakashima, K., Yamaguchi-Shinozaki, K., and Shinozaki, K. (2014). The transcriptional regulatory network in the drought response and its crosstalk in abiotic stress responses including drought, cold, and heat. Front. Plant Sci. 5: 170.

Nolan, T.M., Vukašinović, N., Liu, D., Russinova, E., and Yin, Y. (2020). Brassinosteroids: Multidimensional Regulators of Plant Growth, Development, and Stress Responses. Plant Cell 32: 295–318.

Oka, R., Zicola, J., Weber, B., Anderson, S.N., Hodgman, C., Gent, J.I., Wesselink, J.-J., Springer, N.M., Hoefsloot, H.C.J., Turck, F., and Stam, M. (2017). Genome-wide mapping of transcriptional enhancer candidates using DNA and chromatin features in maize. Genome Biol. 18: 137.

Olvera-Carrillo, Y., Reyes, J.L., and Covarrubias, A.A. (2011). Late embryogenesis abundant proteins. Plant Signaling & Behavior 6: 586–589.

Ordonio, R.L., Ito, Y., Hatakeyama, A., Ohmae-Shinohara, K., Kasuga, S., Tokunaga, T., Mizuno, H., Kitano, H., Matsuoka, M., and Sazuka, T. (2014). Gibberellin deficiency pleiotropically induces culm bending in sorghum: an insight into sorghum semi-dwarf breeding. Sci. Rep. 4: 5287.

Pardo, J., Man Wai, C., Chay, H., Madden, C.F., Hilhorst, H.W.M., Farrant, J.M., and VanBuren, R. (2020). Intertwined signatures of desiccation and drought tolerance in grasses. Proc. Natl. Acad. Sci. U. S. A. 117: 10079–10088.

Parvathaneni, R.K., Bertolini, E., Shamimuzzaman, M., Vera, D., Lung, P.-Y., Rice, B.R., Zhang, J., Brown, P.J., Lipka, A.E., Bass, H.W., and Eveland, A.L. (2020) The regulatory landscape of early maize inflorescence development. Genome Biol. 21: 165.

Qian, D. et al. (2019). Arabidopsis ADF5 promotes stomatal closure by regulating actin cytoskeleton remodeling in response to ABA and drought stress. J. Exp. Bot. 70: 435–446.

Qin, X. and Zeevaart, J.A. (1999). The 9-cis-epoxycarotenoid cleavage reaction is the key regulatory step of abscisic acid biosynthesis in water-stressed bean. Proc. Natl. Acad. Sci. U. S. A. 96: 15354–15361.

Ramírez, F., Dündar, F., Diehl, S., Grüning, B.A., and Manke, T. (2014). deepTools: a flexible platform for exploring deep-sequencing data. Nucleic Acids Res. 42: W187–91.

Ray, D.K., Ramankutty, N., Mueller, N.D., West, P.C., and Foley, J.A. (2012). Recent patterns of crop yield growth and stagnation. Nat. Commun. 3: 1293.

Rentería, M.E., Cortes, A., and Medland, S.E. (2013). Using PLINK for Genome-Wide Association Studies (GWAS) and data analysis. Methods Mol. Biol. 1019: 193–213.

Reynolds, M.P. et al. (2016). An integrated approach to maintaining cereal productivity under climate change. Global Food Security 8: 9–18.

Reynoso, M.A. et al. (2019). Evolutionary flexibility in flooding response circuitry in angiosperms. Science 365: 1291–1295.

Riboni, M., Robustelli Test, A., Galbiati, M., Tonelli, C., and Conti, L. (2016). ABA-dependent control of GIGANTEA signalling enables drought escape via up-regulation of FLOWERING LOCUS T in Arabidopsis thaliana. J. Exp. Bot. 67: 6309–6322.

Rodgers-Melnick, E., Vera, D.L., Bass, H.W., and Buckler, E.S. (2016). Open chromatin reveals the functional maize genome. Proc. Natl. Acad. Sci. U. S. A. 113: E3177–84.

Sah, S.K., Reddy, K.R., and Li, J. (2016). Abscisic Acid and Abiotic Stress Tolerance in Crop Plants. Front. Plant Sci. 7: 571.

Shen, Q., Zhang, P., and Ho, T.H. (1996). Modular nature of abscisic acid (ABA) response complexes: composite promoter units that are necessary and sufficient for ABA induction of gene expression in barley. Plant Cell 8: 1107–1119.

Shi, J., Habben, J.E., Archibald, R.L., Drummond, B.J., Chamberlin, M.A., Williams, R.W., Lafitte, H.R., and Weers, B.P. (2015). Overexpression of ARGOS Genes Modifies Plant Sensitivity to Ethylene, Leading to Improved Drought Tolerance in Both Arabidopsis and Maize. Plant Physiol. 169: 266–282.

Song, L., Huang, S.-S.C., Wise, A., Castanon, R., Nery, J.R., Chen, H., Watanabe, M., Thomas, J., Bar-Joseph, Z., and Ecker, J.R. (2016). A transcription factor hierarchy defines an environmental stress response network. Science 354.

Srivastava, M., Srivastava, A.K., Orosa-Puente, B., Campanaro, A., Zhang, C., and Sadanandom, A. (2020). SUMO Conjugation to BZR1 Enables Brassinosteroid Signaling to Integrate Environmental Cues to Shape Plant Growth. Curr. Biol. 30: 1410–1423.e3.

Stocker, T.F., Qin, D., Plattner, G.-K., Tignor, M., Allen, S.K., Boschung, J., Nauels, A., Xia, Y., Bex, V., Midgley, P.M., and Others (2013). Climate change 2013: The physical science basis. Contribution of working group I to the fifth assessment report of the intergovernmental panel on climate change 1535.

Sullivan, A.M. et al. (2014). Mapping and dynamics of regulatory DNA and transcription factor networks in A. thaliana. Cell Rep. 8: 2015–2030.

Sullivan, A.M. et al. (2019). Mapping and Dynamics of Regulatory DNA in Maturing Arabidopsis thaliana Siliques. Front. Plant Sci. 10: 1434.

Swift, J. and Coruzzi, G.M. (2017). A matter of time - How transient transcription factor interactions create dynamic gene regulatory networks. Biochim. Biophys. Acta Gene Regul. Mech. 1860: 75–83.

Takasaki, H., Maruyama, K., Takahashi, F., Fujita, M., Yoshida, T., Nakashima, K., Myouga, F., Toyooka, K., Yamaguchi-Shinozaki, K., and Shinozaki, K. (2015). SNAC-As, stress-responsive NAC transcription factors, mediate ABA-inducible leaf senescence. Plant J. 84: 1114–1123.

Tuinstra, M.R., Grote, E.M., Goldsbrough, P.B., and Ejeta, G. (1997). Genetic analysis of post-flowering drought tolerance and components of grain development in Sorghum bicolor (L.) Moench. Mol. Breed. 3: 439–448.

Varshney, R.K., Singh, V.K., Kumar, A., Powell, W., and Sorrells, M.E. (2018a). Can genomics deliver climate-change ready crops? Curr. Opin. Plant Biol. 45: 205–211.

Varshney, R.K., Tuberosa, R., and Tardieu, F. (2018b). Progress in understanding drought tolerance: from alleles to cropping systems. Journal of Experimental Botany 69: 3175–3179.

Vera, D.L., Madzima, T.F., Labonne, J.D., Alam, M.P., Hoffman, G.G., Girimurugan, S.B., Zhang, J., McGinnis, K.M., Dennis, J.H., and Bass, H.W. (2014). Differential nuclease sensitivity profiling of chromatin reveals biochemical footprints coupled to gene expression and functional DNA elements in maize. Plant Cell 26: 3883–3893.

Vietmeyer, N.D., Ruskin, F.R., Council, N.R., and Others (1996). Lost crops of Africa. v. 1: Grains.

Wang, H., Tang, J., Liu, J., Hu, J., Liu, J., Chen, Y., Cai, Z., and Wang, X. (2018). Abscisic Acid Signaling Inhibits Brassinosteroid Signaling through Dampening the Dephosphorylation of BIN2 by ABI1 and ABI2. Mol. Plant 11: 315–325.

Wang, L. et al. (2014). Comparative analyses of C₄ and C₃ photosynthesis in developing leaves of maize and rice. Nat. Biotechnol. 32: 1158–1165.

Wang, X., Guo, C., Peng, J., Li, C., Wan, F., Zhang, S., Zhou, Y., Yan, Y., Qi, L., Sun, K., and Others (2019). ABRE-BINDING FACTORS play a role in the feedback regulation of ABA signaling by mediating rapid ABA induction of ABA co-receptor genes. New Phytol. 221: 341–355.

Xiao, B., Huang, Y., Tang, N., and Xiong, L. (2007). Over-expression of a LEA gene in rice improves drought resistance under the field conditions. Theor. Appl. Genet. 115: 35–46.

Xu, Y.-H., Liu, R., Yan, L., Liu, Z.-Q., Jiang, S.-C., Shen, Y.-Y., Wang, X.-F., and Zhang, D.-P. (2012). Light-harvesting chlorophyll a/b-binding proteins are required for stomatal response to abscisic acid in Arabidopsis. J. Exp. Bot. 63: 1095–1106.

Yang, Y. et al. (2020). ABF3 enhances drought tolerance via promoting ABA-induced stomatal closure by directly regulating ADF5 in Populus euphratica. J. Exp. Bot. 71: 7270–7285.

Yoshida, T., Fujita, Y., Sayama, H., Kidokoro, S., Maruyama, K., Mizoi, J., Shinozaki, K., and Yamaguchi-Shinozaki, K. (2010). AREB1, AREB2, and ABF3 are master transcription factors that cooperatively regulate ABRE-dependent ABA signaling involved in drought stress tolerance and require ABA for full activation. Plant J. 61: 672–685.

Yoshida, T., Mogami, J., and Yamaguchi-Shinozaki, K. (2015). Omics Approaches Toward Defining the Comprehensive Abscisic Acid Signaling Network in Plants. Plant Cell Physiol. 56: 1043–1052.

Yu, G., Wang, L.-G., Han, Y., and He, Q.-Y. (2012). clusterProfiler: an R package for comparing biological themes among gene clusters. OMICS 16: 284–287.

Zhao, H., Zhang, W., Zhang, T., Lin, Y., Hu, Y., Fang, C., and Jiang, J. (2020). Genome-wide MNase hypersensitivity assay unveils distinct classes of open chromatin associated with H3K27me3 and DNA methylation in Arabidopsis thaliana. Genome Biol. 21: 24.

