## Supplemental Figures and Tables for "Regulatory signatures of drought response in stress resilient *Sorghum bicolor*"

**Supplemental Figure 1.** Nuclei isolation, nucleosome phasing, and ATAC-seq library QC……………..2

**Supplemental Figure 2.** RNA-seq data showed strong correlations among biological replicates and separation of tissue types by PCA……………………………………………………………………….3

**Supplemental Figure 3.** Up-regulated genes showed enhanced chromatin accessibility around their proximal promoters in response to drought stress…………………………………………………………4

**Supplemental Figure 4.** WGCNA module identification and relationship……………………………….5

**Supplemental Figure 5.** WGCNA module eigengene (ME) expression plots for all modules…………...6

**Supplemental Figure 6.** Plots showing cumulative water usage across the BAP in WW and WS controlled environment conditions………………………………………………………………………7

**Supplemental Figure 7.** Heatmap of WUE residuals over the drought response period for genotypes in subpops 2 and 3 from controlled environment phenotyping experiment…………………………………..8

**Supplemental Figure 8.** Presence/Absence Variation (PAV) in ABF target genes………………………9

**Supplemental Figure 9.** SNP variation within ABF target genes……………………………………….10

**Supplemental Figure 10.** SNP variation within ABRE motifs in DARs………………………………..11

**Supplemental Tables:**

**Supplemental Table 1.** Mapping statistics for ATAC-seq data from inner leaf and crown root………..12

**Supplemental Table 2.** Homer peak call statistics based on ATAC-seq data for the inner leaf and crown root samples…………………………………………………………………………………………13

**Supplemental Table 3.** Distribution of high confidence ATAC-seq peaks across genomic features…..14

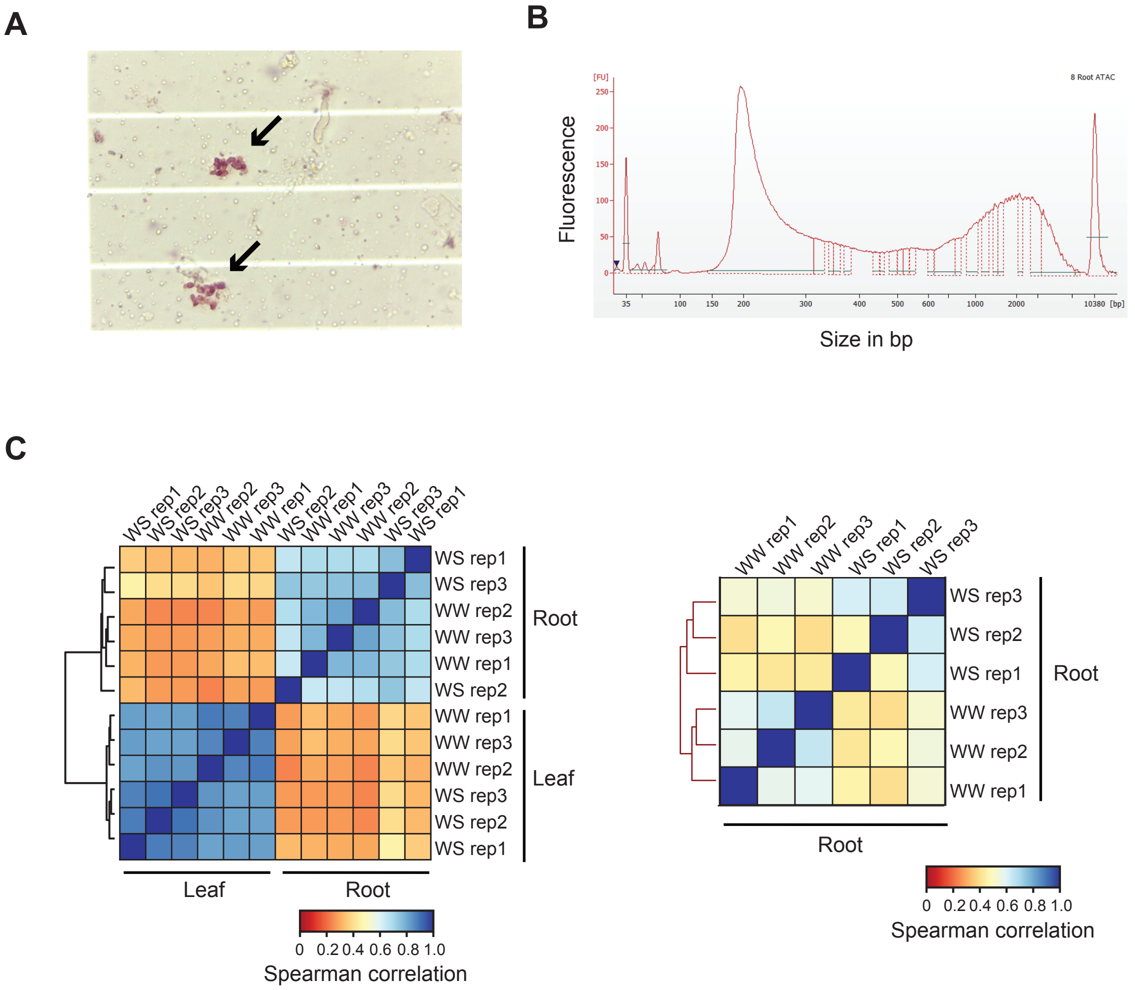


**Supplemental Figure 1. Nuclei isolation, nucleosome phasing, and ATAC-seq library QC. A.** Image of nuclei isolated from leaf tissue viewed in a single haemocytometer cell, which was used to count nuclei for library standardization. Nuclei were stained with acetocarmine dye and visualized in a 40X magnification to check integrity. **B.** A bioanalyzer trace of a representative ATAC-seq library from a root sample. A prominent peak is observed at 200bp (mono-nucleosome plus adapters) with smaller peaks observed at 350bp and 500bp indicative of di- and tri- nucleosome banding, respectively. **C.** Spearman correlations among ATAC-seq libraries between treatments, tissue types, and across biological replicates. Read coverage within all high-confidence peaks was used for clustering. Biological replicates showed strong concordance among the inner leaf samples and slightly lower in root; one lower coverage root WS sample clustered independently when considering correlation of all samples, but clustered with biological replicates when considering only root samples (shown in small heatmap to the right).

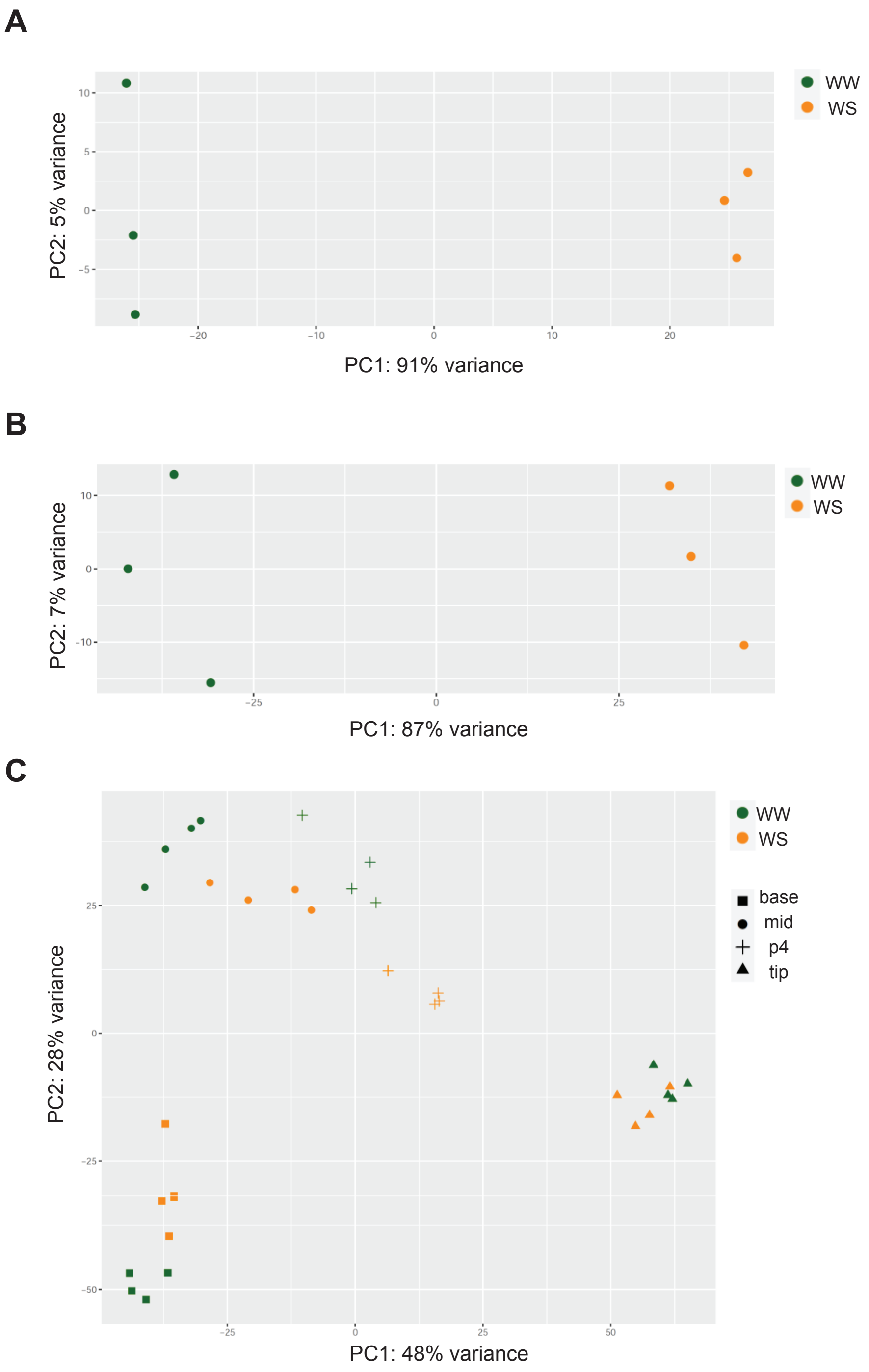


**Supplemental Figure 2. RNA-seq data showed strong correlations among biological replicates and separation of tissue types by PCA.** Principal Components Analysis (PCA) for RNA-seq samples from **A.** inner leaf, **B.** root, and **C.** the leaf gradient tissues.

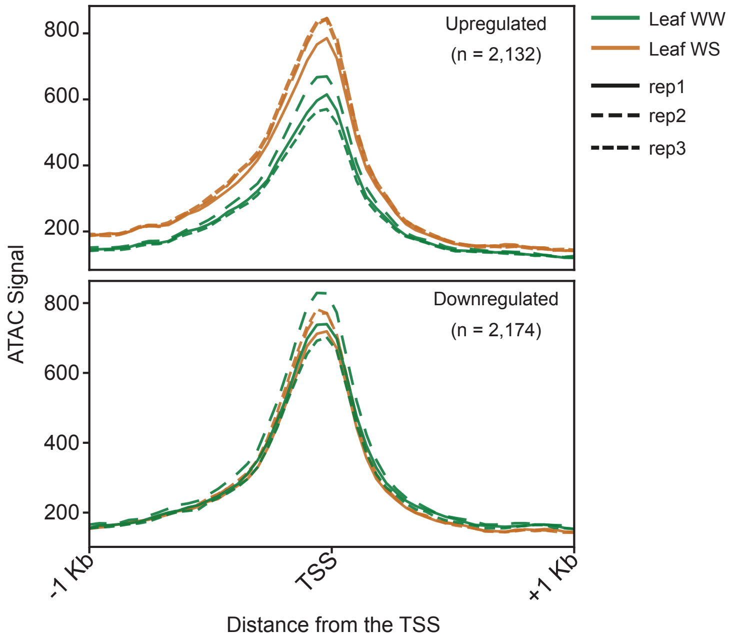


**Supplemental Figure 3: Relationship between accessible chromatin and DE genes.** DE genes were classified based on their up- or down-regulation in response to drought. For each of these genes, the ATAC signal was plotted around the transcriptional start site (TSS) using deeptools plotProfile. The ATAC signal was markedly increased at and proximal to the TSS in up-regulated genes in response to drought.

A

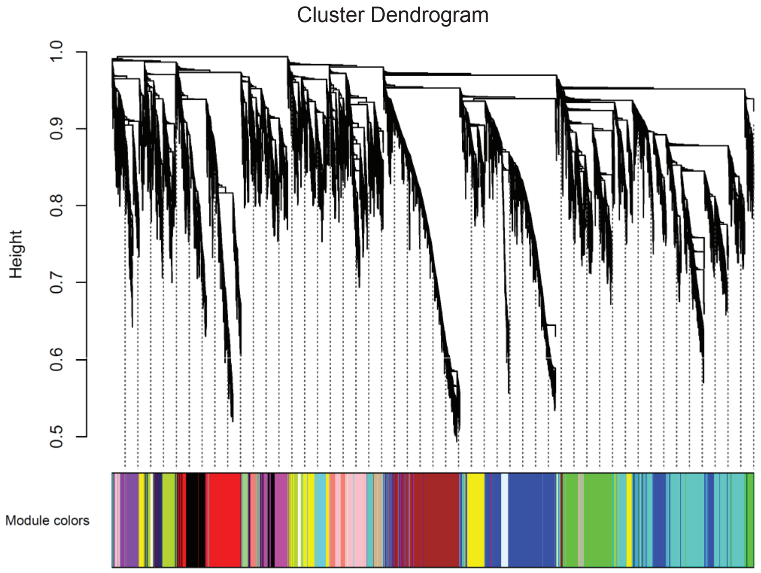


B

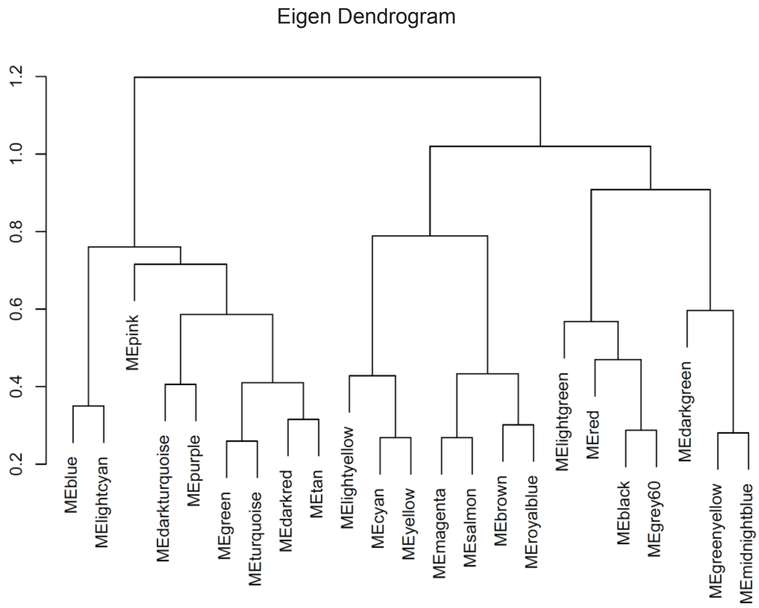


**Supplemental Figure 4. WGCNA module identification and relationship**. **A.** Dendrogram used for dynamic clustering of 23 modules. **B.** Eigengene dendrogram showing the relatedness of modules. Note the close proximity of three of the drought modules (MElightyellow, MEcyan and MEyellow).

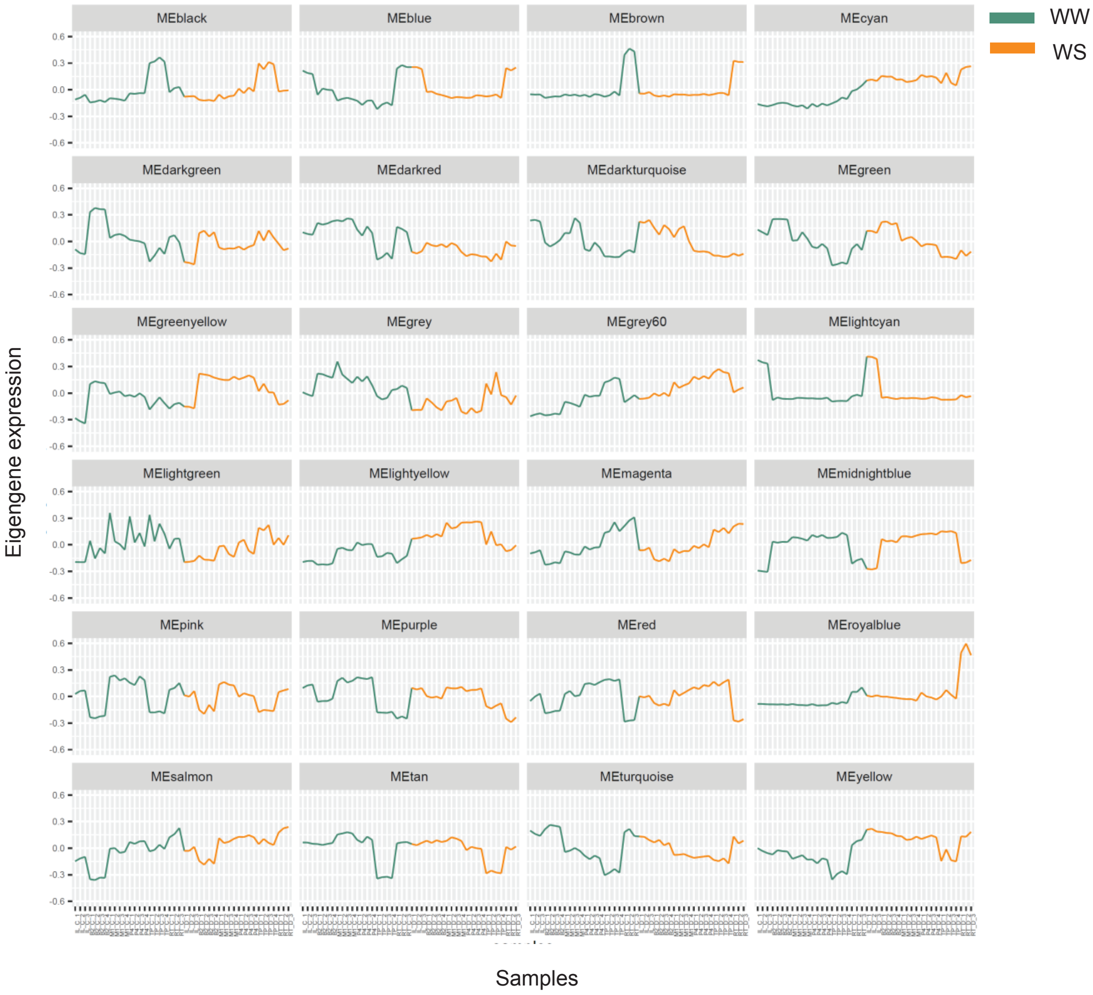


**Supplemental Figure 5. WGCNA module eigengene (ME) expression plots for all modules.** ME can be considered as the most representative gene expression profile of the module and this is shown for all modules with WW and WS samples indicated.

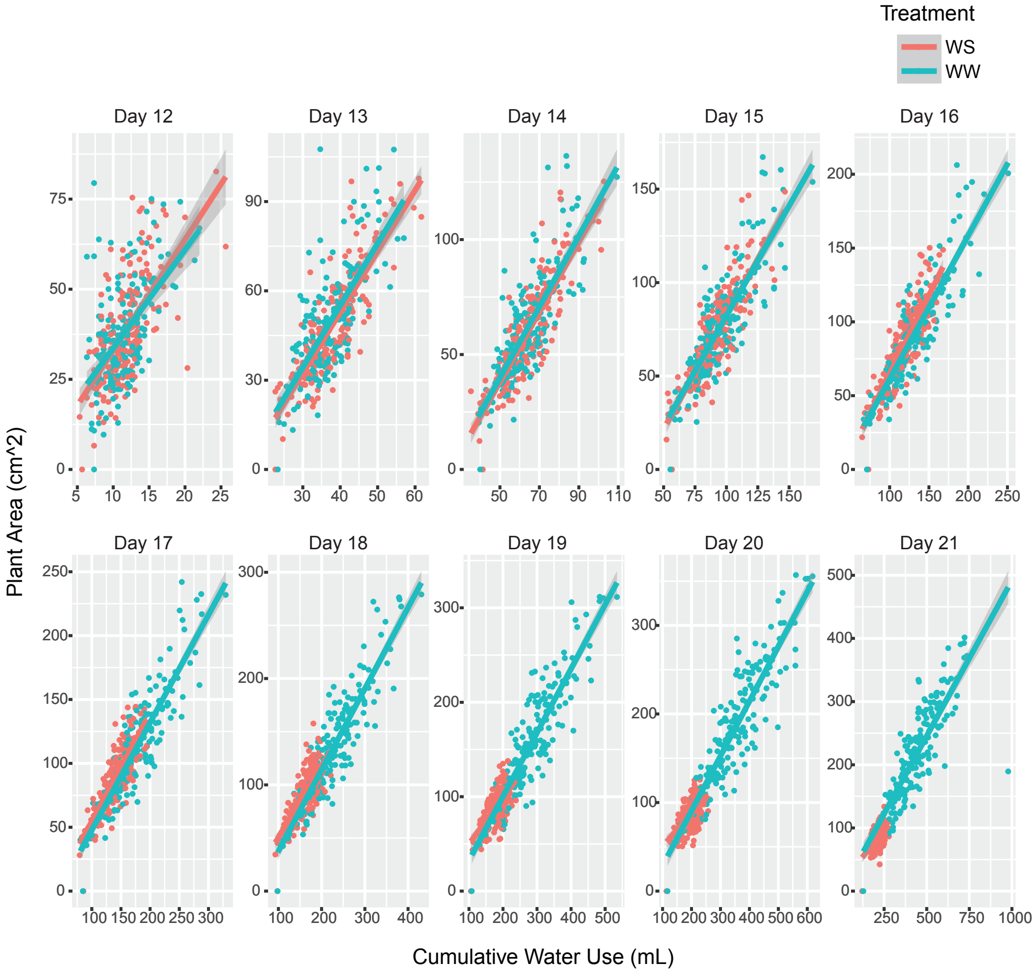


**Supplemental Figure 6. Plots showing cumulative water usage across the BAP in WW and WS controlled environment conditions**. Cumulative water use and plant area measured for each plant on the LemnaTec Scanalyzer platform over a 10 day time frame (Days 12-21) was used to calculate WUE. Plant area was determined using PlantCV (see Methods). WS plants showed lower cumulative water use over time and smaller plant area as compared to WW plants.

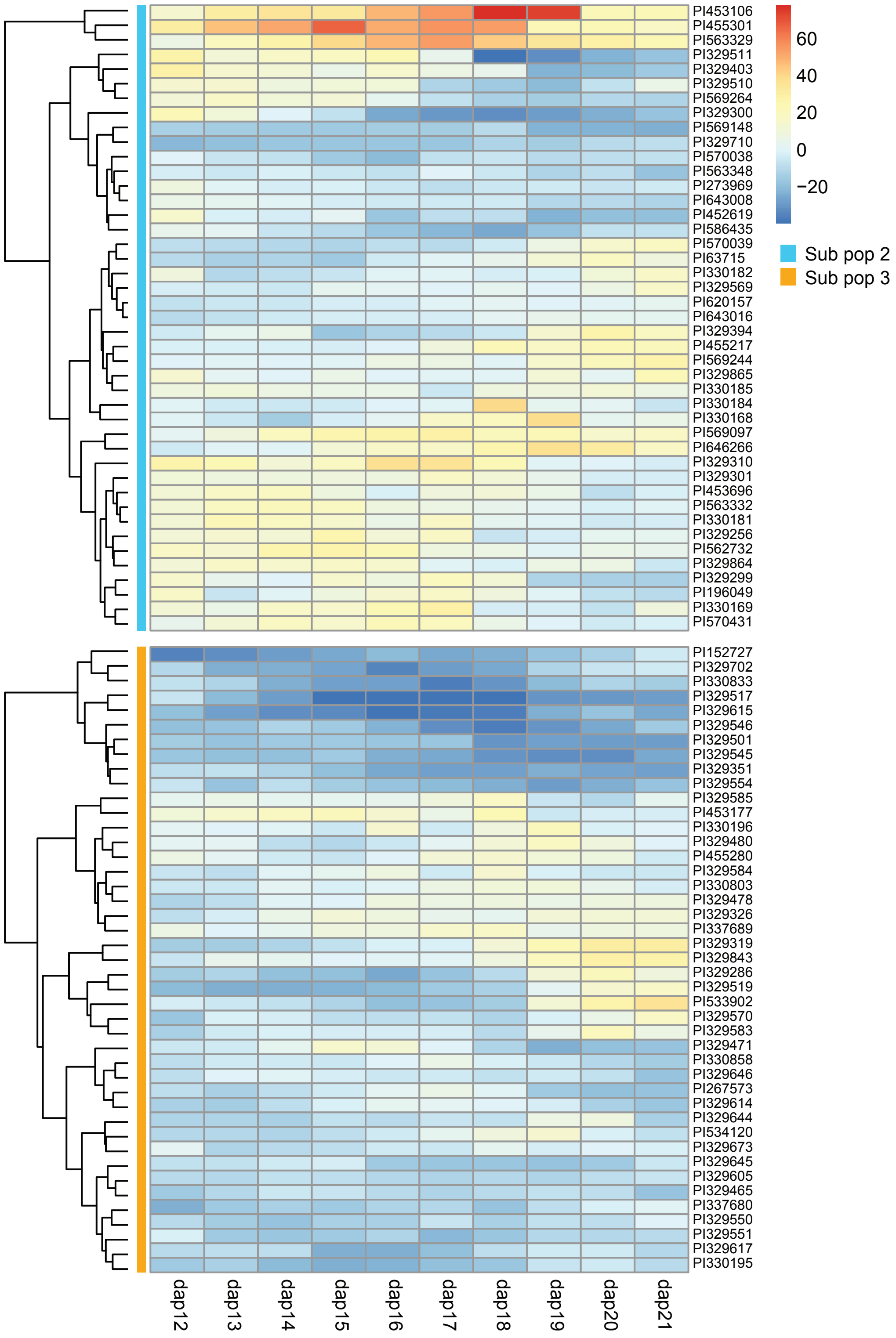


**Supplemental Figure 7. Heatmap of WUE residuals over the drought response period for genotypes in subpops 2 and 3 from controlled environment phenotyping experiment.** Figure 7A showed mean WUE over time was highest for genotypes in subpop 2 and lowest in suppop 3. These data support that cumulative result by showing dynamic WUE of individual genotypes from these subpopulations over time.

A

B

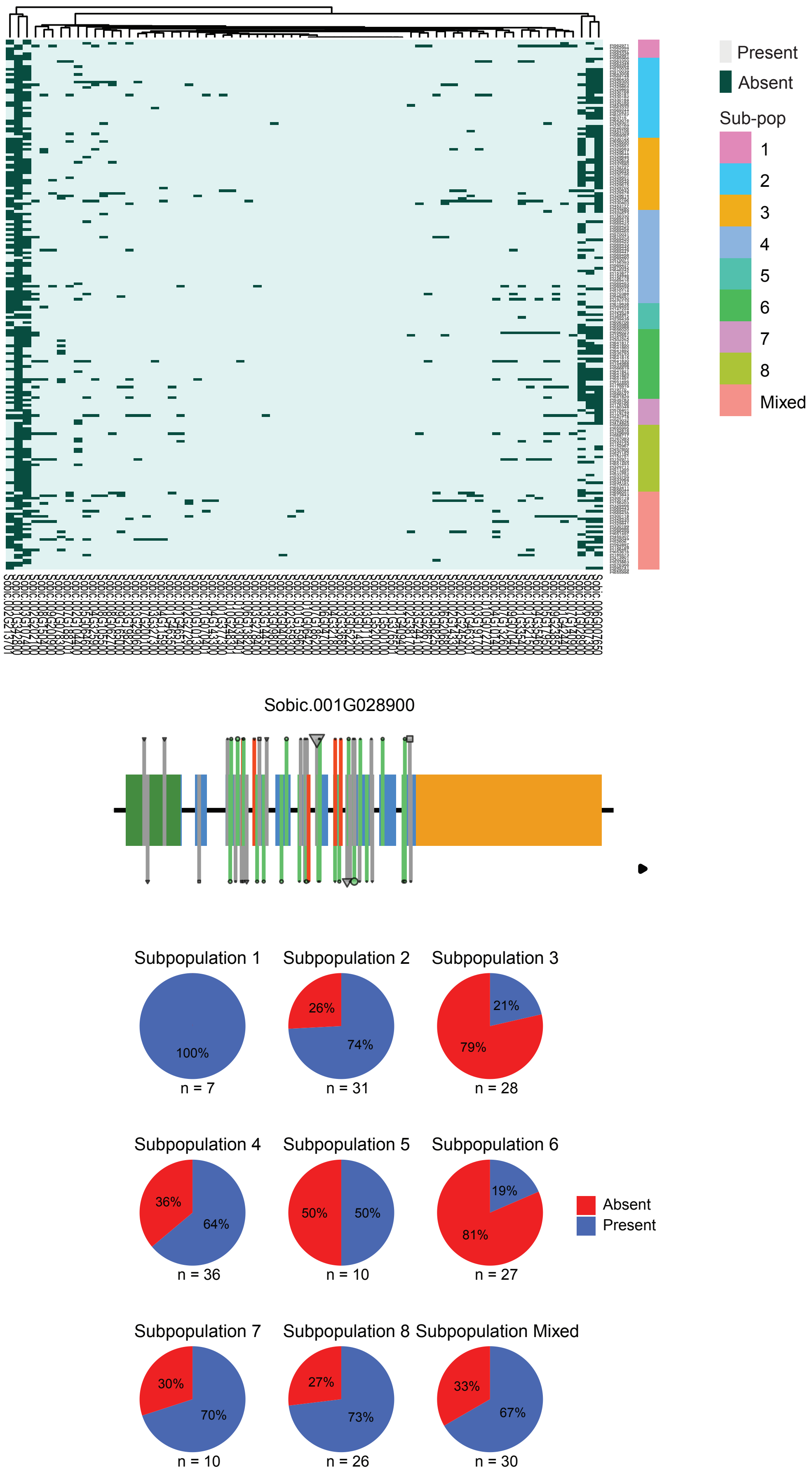


**Supplemental Figure 8. Presence/Absence Variation (PAV) in ABF target genes. A.** Heatmap showing PAV in the 70 predicted target genes of the four ABF TFs based on our network analyses. **B.** The SNP and PAV profile of Sobic.001G028900 (ULP1a).

**A**

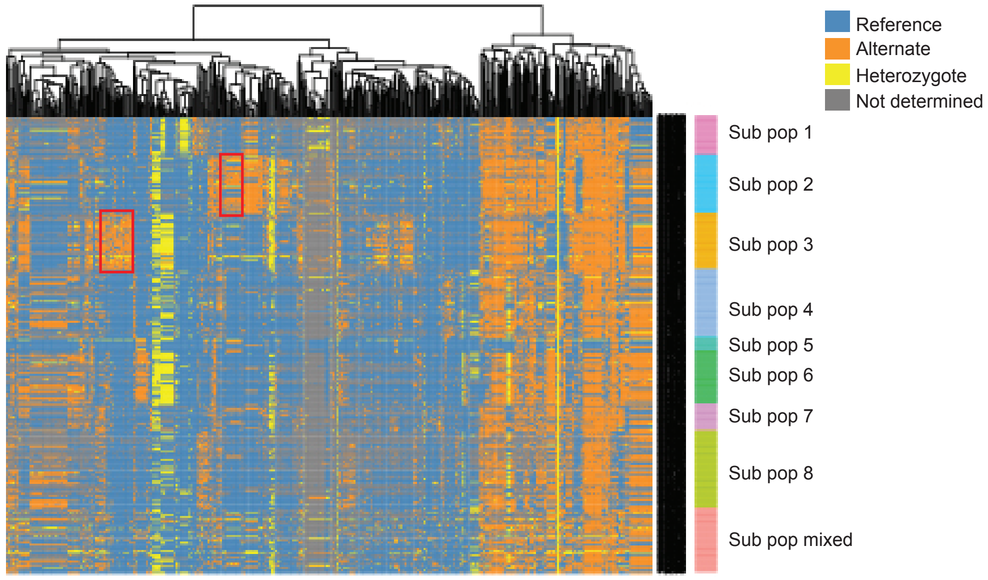


**B**

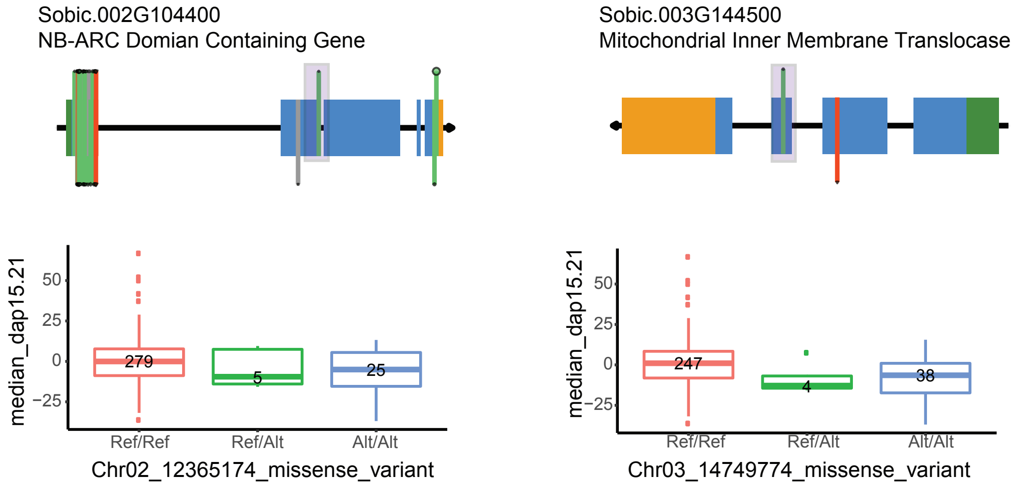


**Supplemental Figure 9. SNP variation within ABF target genes. A.** Heatmap based on supervised clustering shows differences in allele frequencies for SNPs within ABF target genes across subpopulations. A cluster of SNPs that were largely exclusive to subpops 2 or 3 are indicated in the red box. Permutation tests of significance were conducted on these SNPs. **B.** In addition to the WUE-associated SNP in SbGI (Figure 7B), two other ABF target genes harbored SNPs with significant associations for decreased WUE; those encoding a NB-ARC domain protein and a mitochondrial inner membrane translocase. These SNPs are highlighted in the grey boxes.

A

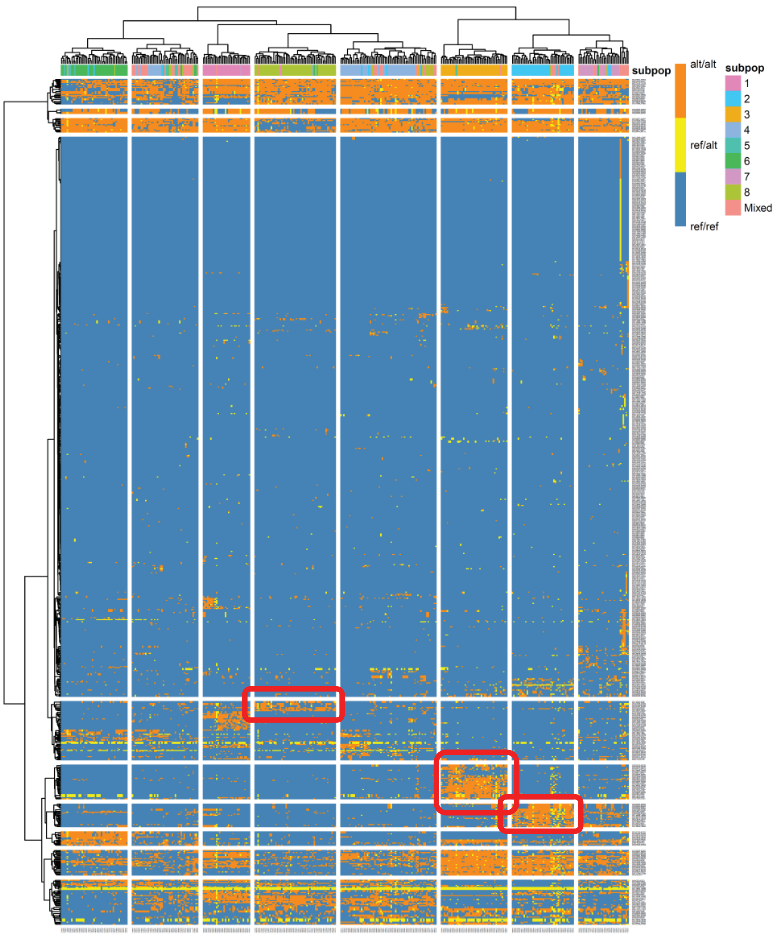


B

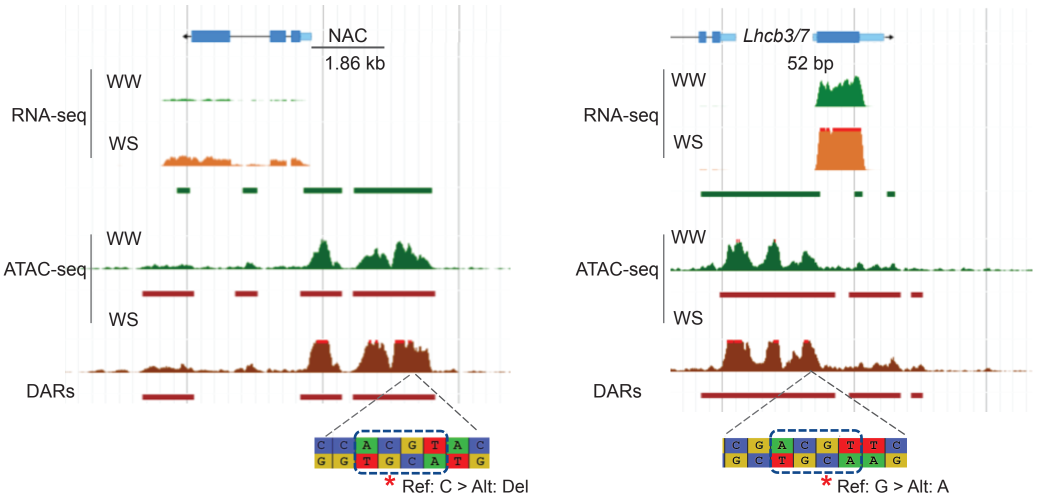


**Supplemental Figure 10. SNP variation within ABRE motifs in DARs. A.** SNP variation within ABRE elements within DARs were clustered based on allele frequencies. Sorghum genotypes are shown along the x-axis and SNPs along the y-axis. Clustering of this specific SNP set separated lines out by subpopulation. Red boxes highlight the subpop dominant SNPs used for significance testing (Supplemental Data Set 15). **B.** Browser views show locations of two subpop 3 dominant polymorphisms within ABRE motifs (red stars) that significantly associated with deceased WUE and were present in the promoter of a gene encoding a NAC TF (Sobic.005G064600; p = 0.026) and Lhcb3/7 (Sobic.009G234600; p = 0.039), which potentially disrupt the core binding motif (boxed).

**Supplemental Table 1. Mapping statistics for ATAC-seq data from inner leaf and crown root.**

[1]Mapped reads to the sorghum genome v 3.1.1 and plastome (chloroplast and mitochondria)

[2]After removal of plastome reads and reads with map quality (MAPQ) < 20

[3]After removal of duplicate reads using picard (see Methods)

| Treatment | Replicate | Total sequenced reads | Total mapped reads[1] | Quality control[2] | Duplicate removal[3] | % of Total mapped reads |
|---|---|---|---|---|---|---|
| *Inner leaf* | | | | | | |
| Control | 1 | 98,883,346 | 96,295,151 | 41,512,555 | 23,250,811 | 24.15 |
| | 2 | 58,431,812 | 57,198,776 | 22,689,849 | 19,954,534 | 34.89 |
| | 3 | 47,029,614 | 46,201,161 | 17,115,456 | 15,536,393 | 33.63 |
| Drought | 1 | 62,136,994 | 61,181,631 | 21,296,420 | 19,129,548 | 31.27 |
| | 2 | 62,434,850 | 60,662,980 | 22,945,029 | 21,130,406 | 34.83 |
| | 3 | 66,060,786 | 64,955,947 | 21,191,937 | 19,118,362 | 29.43 |
| *Root* | | | | | | |
| Control | 1 | 246,772,918 | 55,705,917 | 29,226,845 | 14,289,568 | 25.65 |
| | 2 | 116,109,890 | 28,986,818 | 14,420,903 | 8,706,040 | 30.03 |
| | 3 | 209,992,876 | 56,187,709 | 28,870,545 | 21,017,011 | 37.40 |
| Drought | 1 | 244,981,206 | 65,669,391 | 30,589,392 | 18,379,665 | 27.99 |
| | 2 | 171,133,494 | 53,576,472 | 29,001,804 | 17,573,683 | 32.80 |
| | 3 | 201,541,426 | 67,157,445 | 36,444,569 | 15,891,655 | 23.66 |

**Supplemental Table 2. Homer peak call statistics based on ATAC-seq data for the inner leaf and crown root samples.**

[1]High confidence (HC) peaks were called in peaks were present in at least 2 out of 3 replicates in each treatment (minimum overlap 25%)
[2]HC peaks as a percentage of total peaks in each replicate
[3]HC peaks of each treatment were merged to generate a set of unique peak regions consistent among biological replicates.

| Treatment | Replicate | Peak # | Median size | HC peaks[1] | % Total peaks[2] | Merged HC peaks[3] |
|---|---|---|---|---|---|---|
| *Inner leaf* | | | | | | |
| Drought | 1 | 70,660 | 324 | 58,853 | 83.29 | 58,654 |
| | 2 | 64,015 | 316 | 59,655 | 93.19 | |
| | 3 | 57,946 | 304 | 56,335 | 97.22 | |
| Control | 1 | 62,765 | 312 | 57,275 | 91.25 | 57,518 |
| | 2 | 64,775 | 317 | 57,570 | 88.88 | |
| | 3 | 61,701 | 306 | 56,289 | 91.23 | |
| *Root* | | | | | | |
| Drought | 1 | 31,483 | 243 | 28413 | 90.25 | 29951 |
| | 2 | 24,222 | 230 | 23196 | 95.76 | |
| | 3 | 53,941 | 317 | 27193 | 50.41 | |
| Control | 1 | 37,738 | 263 | 32011 | 84.82 | 35,835 |
| | 2 | 45,085 | 277 | 34813 | 77.22 | |
| | 3 | 42,648 | 277 | 34861 | 81.74 | |

**Supplemental Table 3. Distribution of the high confidence ATAC-seq peaks across the genomic features.**

[1]In several cases, overlapping gene models can result in assignment of one ATAC-seq peak to multiple features
[2]For genomic distribution analyses, the control high confidence (HC) peaks were used from inner leaf (n=57,518) and root (n = 35,835) and the DARs (inner-leaf (n =6,496) and root (n=2,275))
[3]The promoter is considered 2 kb upstream of the transcription start site (TSS)
[4]The proximal promoter is 1kb upstream and 200bp downstream of the TSS
[5]The HC peaks were normalized to a factor of 10kb and DARs were normalized to a factor of 1Mb
[6]The UTRs overlapping with the exon annotations were removed

| Genomic feature[1] | Size of the feature (in bp) | Inner leaf[2] | % of total peaks | Leaf normalized[5] | Root[2] | % of Total peaks | Root normalized |
|---|---|---|---|---|---|---|---|
| HC peaks | | | | | | | |
| Promoter[3] | 52,932,111 | 15,878 | 27.61 | 3.00 | 11,749 | 32.79 | 2.22 |
| Proximal Promoter[4] | 39,540,408 | 19,323 | 33.59 | 4.89 | 14,545 | 40.59 | 3.68 |
| 5' UTR | 8,404,213 | 7,526 | 13.08 | 8.96 | 5,556 | 15.5 | 6.61 |
| Exon[6] | 39,408,245 | 4,887 | 8.5 | 1.24 | 2,432 | 6.79 | 0.62 |
| Intron | 62,166,720 | 4,373 | 7.6 | 0.70 | 2,614 | 7.29 | 0.42 |
| 3' UTR | 14,385,383 | 3,410 | 5.93 | 2.37 | 1,831 | 5.11 | 1.27 |
| Flank 1K | 25,640,478 | 6,293 | 10.94 | 2.45 | 3,851 | 10.75 | 1.50 |
| Intergenic | 488,799,404 | 16,965 | 29.5 | 0.35 | 9,032 | 25.2 | 0.18 |
| DAR distribution | | | | | | | |
| Promoter[3] | 52,932,111 | 1,584 | 24.38 | 2.99 | 632 | 27.78 | 1.19 |
| Proximal Promoter[4] | 39,540,408 | 1,632 | 25.12 | 4.13 | 646 | 28.40 | 1.63 |
| 5' UTR | 8,404,213 | 520 | 8.00 | 6.19 | 196 | 8.62 | 2.33 |
| Exon | 39,408,245 | 575 | 8.85 | 1.46 | 199 | 8.75 | 0.50 |
| Intron | 62,166,720 | 583 | 8.97 | 0.94 | 217 | 9.54 | 0.35 |
| 3' UTR | 14,385,383 | 422 | 6.50 | 2.93 | 192 | 8.44 | 1.33 |
| Flank 1K | 25,640,478 | 674 | 10.38 | 2.63 | 247 | 10.86 | 0.96 |
| Intergenic | 488,799,404 | 2,318 | 35.68 | 0.47 | 664 | 29.19 | 0.14 |
